# Machine-learning analysis of factors that shape cancer aneuploidy landscapes reveals an important role for negative selection

**DOI:** 10.1101/2023.07.05.547626

**Authors:** Juman Jubran, Rachel Slutsky, Nir Rozenblum, Lior Rokach, Uri Ben-David, Esti Yeger-Lotem

**Affiliations:** Department of Clinical Biochemistry and Pharmacology, Ben-Gurion University of the Negev, Beer Sheva 84105, Israel; Department of Human Molecular Genetics and Biochemistry, Faculty of Medicine, Tel Aviv University, Tel Aviv, Israel; Department of Software & Information Systems Engineering, Ben-Gurion University of the Negev, Beer Sheva 84105, Israel; The National Institute for Biotechnology in the Negev, Ben-Gurion University of the Negev, Beer Sheva 84105, Israel

## Abstract

Aneuploidy, an abnormal number of chromosomes within a cell, is considered a hallmark of cancer. Patterns of aneuploidy differ across cancers, yet are similar in cancers affecting closely-related tissues. The selection pressures underlying aneuploidy patterns are not fully understood, hindering our understanding of cancer development and progression. Here, we applied interpretable machine learning (ML) methods to study tissue-selective aneuploidy patterns. We defined 20 types of features of normal and cancer tissues, and used them to model gains and losses of chromosome-arms in 24 cancer types. In order to reveal the factors that shape the tissue-specific cancer aneuploidy landscapes, we interpreted the ML models by estimating the relative contribution of each feature to the models. While confirming known drivers of positive selection, our quantitative analysis highlighted the importance of negative selection for shaping the aneuploidy landscapes of human cancer. Tumor-suppressor gene density was a better predictor of gain patterns than oncogene density, and vice-versa for loss patterns. We identified the contribution of tissue-selective features and demonstrated them experimentally for chr13q gain in colon cancer. In line with an important role for negative selection in shaping the aneuploidy landscapes, we found compensation by paralogs to be a top predictor of chromosome-arm loss prevalence, and demonstrated this relationship for one such paralog interaction. Similar factors were found to shape aneuploidy patterns in human cancer cell lines, demonstrating their relevance for aneuploidy research. Overall, our quantitative, interpretable ML models improve the understanding of the genomic properties that shape cancer aneuploidy landscapes.

## Introduction

Aneuploidy, defined as an abnormal number of chromosomes or chromosome-arms within a cell, is a characteristic trait of human cancer (Ben-David and Amon 2020). Aneuploidy is associated with patient prognosis and with response to anticancer therapies (Shukla et al. 2020; Vasudevan et al. 2020), indicating that it can play a driving role in tumorigenesis. It is well established that the fitness advantage conferred by specific aneuploidies depends on the genomic, environmental and developmental contexts (Ben-David and Amon 2020). One important cellular context is the cancer tissue-of-origin; aneuploidy patterns are cancer type-specific, and cancers that originate from related tissues tend to exhibit similar aneuploidy patterns (Ben-David et al. 2017; Shukla et al. 2020; Taylor et al. 2018). Nonetheless, the selection pressures that shape the aneuploidy landscapes of human tumors are not fully understood, and it is not clear why some chromosome-arm gains and losses would recur in some tumor types but not in others.

Several non-mutually-exclusive explanations have been previously provided in an attempt to explain the tissue selectivity of aneuploidy patterns. First, the densities of oncogenes (OGs) and tumor suppressor genes (TSGs) are enriched in chromosome-arms that tend to be gained or lost, respectively, potentially due to the cumulative effect of altering multiple such genes at the same time (Davoli et al. 2013). As cell proliferation is controlled in a tissue-dependent manner, the relative importance of OGs and TSGs varies across cell types, so that the density of tissue-specific driver genes can help predict aneuploidy patterns (Sack et al. 2018). Second, the tissue-specific chromosome-wide gene expression profiles also contribute to the observed cancer type-specific aneuploidy patterns, suggesting that chromosome-arm gains and losses may “hardwire” gene expression patterns that are characteristic of the tumor tissue-of-origin (Patkar et al. 2021). Third, several strong cancer driver genes have been shown to underlie the recurrent aneuploidy of the chromosome-arms on which these genes reside; prominent examples are the tumor suppressors *TP53* and *PTEN*, which have been shown to drive the recurrent loss of chromosome-arm 17p in leukemia and that of 10p in glioma, respectively (Liu et al. 2016; Zhou et al. 1999). Fourth, it has been recently proposed that somatic amplifications, including chromosome-arm gains, are positively selected in cancer evolution in order to buffer gene inactivation of haploinsufficient genes in mutation-prone regions (Alfieri, Caravagna, and Schaefer 2023).

Notably, each previous study focused on a separate aspect of tissue-specificity; therefore, the relative contribution of each factor to shaping the overall aneuploidy landscape of human tumors is currently unknown. Furthermore, whether any additional tissue-specific traits could also play a major role in driving aneuploidy patterns, remains an open question. Importantly, previous studies focused on the role of positive selection in driving the gain or the loss of specific chromosome-arms in specific tumor-types. However, unlike point mutations in specific genes, aneuploidies come with a strong fitness cost (Ben-David and Amon 2020; Sheltzer and Amon 2011). Therefore, whereas positive selection greatly outweighs negative selection in shaping the landscape of point mutations in cancer (Martincorena et al. 2017), both positive selection and negative selection may be important for shaping the landscape of aneuploidy. Indeed, a recent preprint showed that negative selection could determine the boundaries of recurrent cancer copy-number alterations (Shih et al. 2023). It is therefore necessary to consider the balance between positive and negative selection in shaping the aneuploidy landscape of human cancers.

Machine-learning (ML) methods have been applied to study a variety of biological and medical questions where heterogeneous large-scale data are available (Zitnik et al. 2019). In the context of cancer, supervised ML methods were applied to predict cancer driver genes (Han et al. 2019; Luo et al. 2019), to distinguish between cancer types (Mostavi et al. 2021; Ramirez et al. 2020), and to predict gene dependency in tumors (Chiu et al. 2021). However, ML has not been applied to investigate the observed patterns of aneuploidy in human cancer. Whereas ML has been frequently used for prediction and often regarded as a black box, recent advancements have allowed more insight into the factors that drive prediction. For example, Shapley Additive exPlanations algorithm (SHAP) (Lundberg and Lee 2017; Rodriguez-Perez and Bajorath 2020) estimates the importance and relative contribution of each of the features utilized by the model to the model’s decisions.

Here, we present a novel ML approach to elucidate the factors that drive the cancer type-specific patterns of aneuploidy. For this, we constructed separate ML models for chromosome-arm gain and loss, whereby each of 39 chromosome-arms within 24 cancer types was associated with 20 types of features corresponding to various genomic attributes of chromosome-arms, normal tissues, primary tumors, and cancer cell lines (CCLs). Our approach is focused on interpretation rather than prediction of aneuploidy recurrence patterns. Interpretation of the gain and loss models for aneuploidy in primary tumors captured known genomic features that had been previously reported to shape aneuploidy landscapes, supporting the models’ validity. Furthermore, these analyses suggested that negative selection played a greater role than positive selection in this process, and revealed paralog compensation as an important contributor to cancer type-specific aneuploidy patterns, in both primary tumors and CCLs. Lastly, we experimentally validated a specific aneuploidy driver using genetically-engineered isogenic human cells.

## Results

### Constructing machine learning models to classify cancer aneuploidy patterns

To create an ML model that predicts the recurrence pattern of aneuploidy across tumor types, we collected data of aneuploidy patterns of chromosome-arms in different cancer types from The Cancer Genome Atlas (TCGA). We focused on 24 cancer types with available transcriptomic data of normal tissue-of-origins (Consortium 2020) (Methods). We used GISTIC2.0 (Mermel et al. 2011) to label each pair of chromosome-arm and cancer type as recurrently gained (199 pairs), recurrently lost (307 pairs) or neutral (430 pairs) (Fig. 1A).

**Figure 1.**
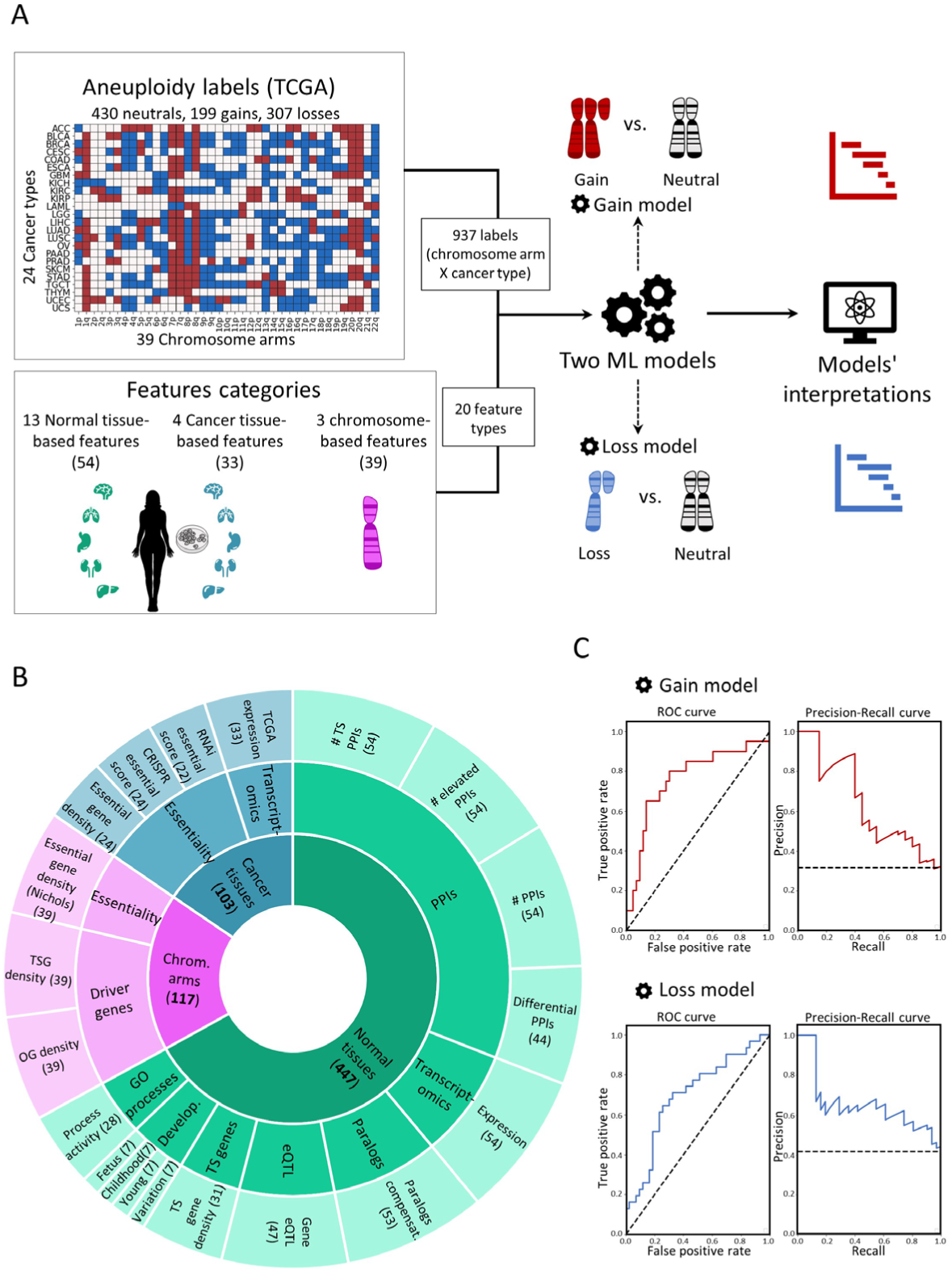
A machine learning (ML) approach for predicting aneuploidy in cancer. A. Schematic view of the ML model construction. Labels represent aneuploidy status of each chromosome arms in 24 cancer types (abbreviation of cancer types detailed in Table S1), classified as gained (red, n = 199), lost (blue, n = 307), or neutral (white, n = 430). Features consist of 20 types of features pertaining to chromosome-arms, normal tissues and cancer tissues (see panel B). Two separate ML models were constructed to predict gained and lost chromosome-arms (gain model and loss model). Each model was analyzed to estimate the contribution of the features to the predicted outcome. B. The features analyzed by the ML model. The inner layer shows feature categories: chromosome arms (purple), cancer tissues (blue), and normal tissues (green). The middle layer shows the sub-categories of the features. Chromosome-arm features include essentiality and driver genes features. Cancer-tissue features include transcriptomics and essentiality features. Normal-tissue features include protein-protein interactions (PPIs), transcriptomics, paralogs, eQTL, tissue-specific (TS) genes, development and GO processes features. The outer layer represents all 20 feature types that were analyzed by the model. Numbers in parentheses indicate the number of tissues, organs, or cell lines from which cancer and normal tissue features were derived, or the number of chromosome-arms from which chromosome-arm features were derived. C. The performance of the ML models as evaluated by the area under the receiver-operating characteristic curve (auROC, left) and the precision recall curve (auPRC, right) using 10-fold cross-validation. Gain model (gradient boosting): auROC=74%, auPRC =63% (expected 32%). Loss model (XGBoost): auROC=70%, auPRC=63% (expected 42%).

Next, we constructed a large-scale dataset consisting of three categories of features (Fig. 1B; Methods). The first category, denoted ‘chromosome-arms’, contained features of chromosome-arms that are independent of cancer type. It included the density of OGs and TSGs (Davoli et al. 2013) and the density of essential genes (Nichols et al. 2020) per chromosome-arm. The second category, denoted ‘cancer tissues’, contained features of pairs of chromosome-arms and cancer tissues, such as the expression levels (TCGA) and essentiality scores (Tsherniak et al. 2017) of genes located on the arm in that cancer tissue, which were inferred from 103 omics-based readouts (Methods). The third category, denoted ‘normal tissues’, contained features pertaining to gene functions and interactions on each chromosome-arm in the normal tissues from which each cancer type originated (Table S1). These were inferred from 447 tissue-based properties. The rich available omics data from normal tissues allowed us to include features that were shown to play a role in other diseases, such as the expression levels of genes located on the arm in the respective normal tissue (Patkar et al. 2021), their tissue protein-protein interactions (PPIs) (Basha et al. 2020; Greene et al. 2015), activity in the tissue of biological processes related to arm genes (Sharon et al. 2022), and dosage relationships between paralogous genes, denoted ‘paralog compensation’ (Barshir et al. 2018; Jubran et al. 2020). Notably, to enhance our understanding of tissue-selectivity, feature values were relative rather than absolute; For example, instead of indicating the absolute expression of genes in a given normal tissue, the feature value was set to the expression of genes in the given tissue relative to all tissues (Methods, Fig. S1, Fig. S2). To fit the features dataset and the labels dataset we further transformed the features dataset, such that each pair of chromosome-arm and cancer type was associated with features corresponding to the chromosome-arm, cancer type, and matching normal tissue (Methods). In total, the dataset included 20 types of features per chromosome-arm and cancer type: three in the chromosome-arm category, four in the cancer tissues category, and thirteen in the normal tissues category (Fig. 1B).

With these labels and features of each chromosome-arm and cancer type, we set out to construct two separate ML models to predict the chromosome-arm gain and loss patterns across cancer types (denoted as the “gain model” and the “loss model”, respectively; Fig. 1A). Each model was trained and tested on data of gained (or lost) chromosome-arms vs. neutral chromosome-arms. We employed five different ML methods, and assessed the performance of each method by using 10-fold cross-validation and calculating average area under the receiver operating characteristic (auROC) and average area under the precision-recall curve (auPRC) (Fig. S3A,B). Logistic regression showed similar results to a random prediction, with auROC of 54% for each model (Fig. S3), indicating that the relationships between features and labels are non-linear. Decision tree methods that can capture such relationships (Kingsford and Salzberg 2008; Kotsiantis 2013), including gradient boosting, XGBoost, and random forest, performed better than logistic regression and similarly to each other (Fig. S3). Best performance in the gain model was achieved by gradient boosting method, with auROC of 74% and auPRC of 63% (expected: 32%) (Fig. 1C). Best performance in the loss model was achieved by XGBoost, with auROC of 70% and auPRC of 63% (expected: 42%) (Fig. 1C). Similar results were obtained upon training and testing models on data of gained (or lost) chromosome-arms against all other chromosome-arms (that is, including not only neutral chromosome-arms but also those altered in the opposite direction; Methods; Fig. S3C,D).

### Revealing the top contributors to cancer aneuploidy patterns

The main purpose of our models was to identify the features that contribute the most to the recurrence patterns of aneuploidy observed in human cancer, which could illuminate the factors at play. To this aim, we used the SHAP (Shapley Additive exPlanations) algorithm (Lundberg and Lee 2017; Rodriguez-Perez and Bajorath 2020), which estimates the importance and relative contribution of each feature to the model‘s decision and ranks them accordingly. We applied SHAP separately to the gain model and to the loss model.

In the gain model, the topmost features were ‘TSG density’ and ‘OG density’ (Fig. 2A,B, Fig. S4). As expected, these features showed opposite directions: TSG density was low in gained chromosome-arms, whereas OG density was high, in line with previous observations (Davoli et al. 2013; Sack et al. 2018) (Fig. 2B). Importantly, this analysis revealed that the impact of TSGs on the gain model‘s decision was twice larger than that of OGs (Fig. 2A), highlighting the importance of negative selection for shaping cancer aneuploidy patterns. The third most important feature was ‘TCGA expression’, which quantified the expression of arm-residing genes in the given cancer type relative to their expression in other cancers. Notably, expression levels were computed independently of arm gain (i.e., excluding samples with the respective chromosome-arm gain; see Methods). This analysis revealed that, across cancer types, chromosome-arms that tend to be gained exhibit higher expression of genes even in neutral cases, consistent with a previous recent study (Patkar et al. 2021). This confirms that the genes on gained chromosome-arms are preferentially important for the specific cancer types in which these gains are recurrent. Congruently, PPIs and normal tissue expression – features of normal tissues – were also among the 10 top-contributing features (Fig. 2A). The estimated importance of all features in the gain model is shown in Fig. S4A.

**Figure 2.**
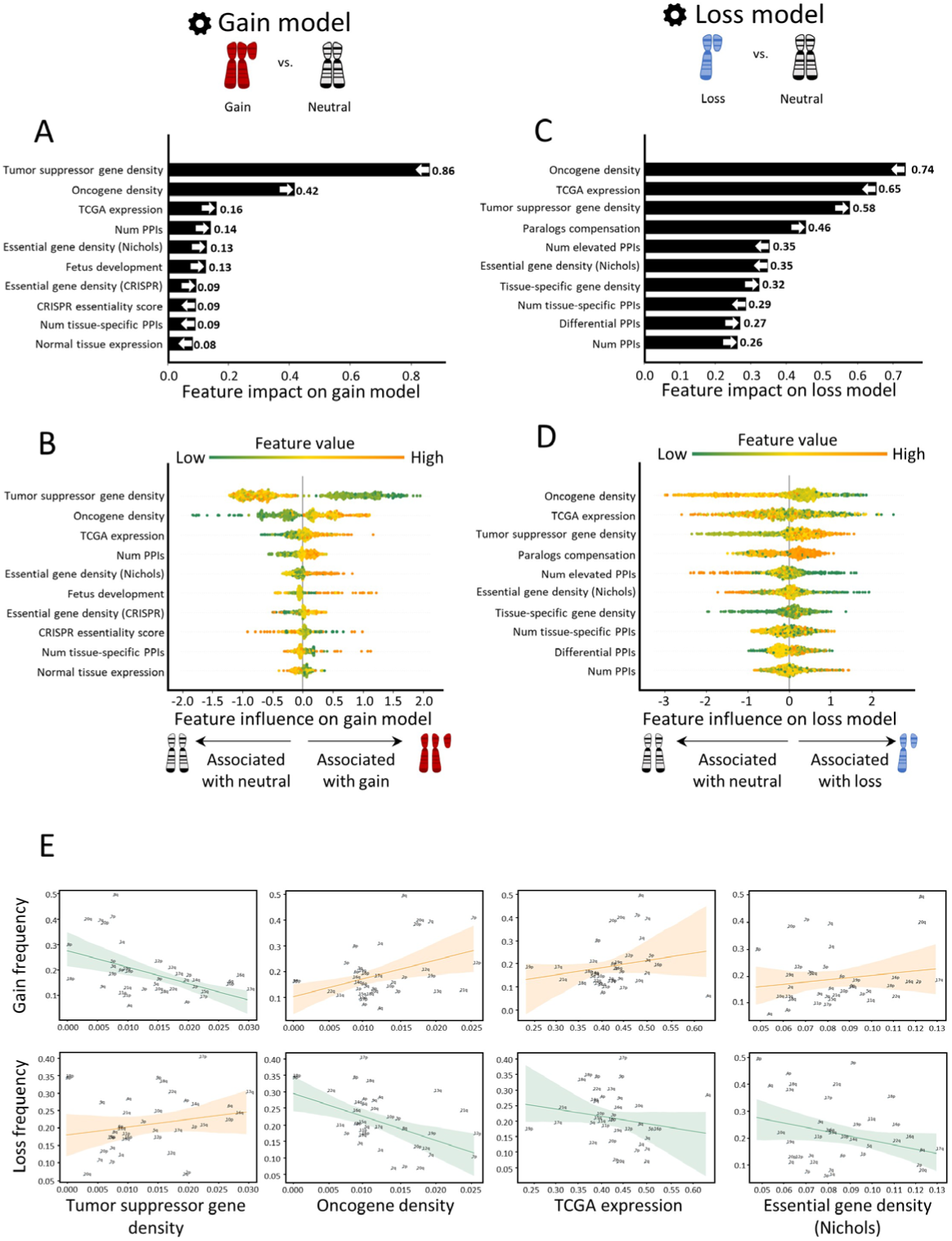
Quantitative views into the 10 topmost contributing features of the gain and loss models. Features are ordered from bottom to top by their increased average absolute contribution on the model. A. The average absolute contribution of each feature to the gain model. The directionality of the feature (i.e., whether high values correspond to gain or neutral) is represented by an arrow. B. A detailed view of the contribution of each feature to the gain model. Per feature, each dot represents the contribution per instance of a chromosome-arm and cancer type pair. The dots are spread based on whether they were classified as neutral (left) or gain (right) by the model. Instances are colored by the feature value (green-to-orange scale denotes low-to-high value). C. Same as panel A for the loss model. D. Same as panel B for the loss model. E. The correlation between top contributing features and the frequencies of chromosome-arms gain and loss, as measured by Spearman correlation. P-values were adjusted for multiple hypothesis testing using Benjamini-Hochberg procedure. Negative correlation between TSG density and gain frequency (ρ=-0.52, adjusted p=0.006). Positive correlation between TSG density and loss frequency (ρ=0.3, adjusted p=0.17). Positive correlation between OG density and gain frequency (ρ=0.25, adjusted p=0.18). Negative correlation between OG density and loss frequency (ρ=-0.47, adjusted p=0.01). Positive correlation between TCGA expression and gain frequency (ρ=0.29, adjusted p=0.14). Negative correlation between TCGA expression and loss frequency (ρ=-0.33, adjusted p=0.12). Positive correlation between essential gene density and gain frequency (ρ=0.16, adjusted p=0.37). Negative correlation between essential gene density and loss frequency (ρ=-0.1, adjusted p=0.5).

The loss model shared the same top three features, yet with opposite directions and different ranks (Fig. 2C,D). ‘OG density’, ranked first, was low in lost chromosome-arms, whereas ‘TSG density’, ranked third, was high (Fig. 2D), in line with previous observations (Davoli et al. 2013; Sack et al. 2018). In contrast to the gain model, in the loss model the impact of OGs on the model‘s decision was larger than that of TSGs, again in line with negative selection as an important force in cancer aneuploidy evolution. ‘TCGA expression’ (computed independently of chromosome-arm loss, see Methods) ranked second: chromosome-arms with highly-expressed genes tended not to be recurrently lost, in line with negative selection. Another top feature that showed opposite directions between the gain and loss model was ‘essential gene density’ (Nichols et al. 2020). As expected, ‘essential gene density’ was low in lost chromosome-arms, in line with negative selection against losing copies of essential genes (McFarland et al. 2018; Nichols et al. 2020; Tsherniak et al. 2017). The estimated importance of all features in the loss model is shown in Fig. S4B.

To examine the direct relationships between high-ranking features and aneuploidy recurrence patterns, we assessed the correlations between these features and aneuploidy prevalence (Methods). In accordance with the SHAP analysis, the negative correlation between ‘TSG density’ and chromosome-arm gain (ρ=-0.52, adjusted p=0.0006, Spearman correlation; Fig. 2E) was much stronger and more significant than the positive correlation between ‘OG density’ and chromosome-arm gain (ρ=0.25, adjusted p=0.12, Spearman correlation; Fig. 2E). Similarly, the negative correlation between ‘OG density’ and chromosome-arm loss (ρ=-0.47, adjusted p=0.003, Spearman correlation; Fig. 2E) was much stronger and more significant than the positive correlation between ‘TSG density’ and chromosome-arm loss (ρ=0.3, adjusted p=0.067, Spearman correlation; Fig. 2E). TCGA expression and essential gene density were correlated with chromosome-arm gain, and anticorrelated with chromosome-arm loss, albeit to a lesser extent (Fig. 2E, Fig. S5). Also showing positive correlations with gains and negative correlations with losses were features derived from expression levels in normal adult and developing tissues, certain PPI-related features, and additional essentiality features (Fig. S5). However, these correlations were weaker than the correlations described above.

Applying the same approaches, we constructed two ML models where we modeled separately gains and losses of chromosome-arm vs. ungained chromosome-arms (i.e., lost and neutral) and unlost chromosome-arms (i.e., gained and neutral), respectively. SHAP analysis of the models revealed that feature importance was very similar between these models and the models comparing gained and lost chromosome-arms to neutral chromosome-arms only (Fig. S6).

### Similar predictive features in human cancer cell lines and in human tumors

Next, we aimed to test whether similar features also shape aneuploidy patterns in CCLs. We collected data of aneuploidy patterns of all chromosome-arms in CCLs (Cohen-Sharir et al. 2021), and analyzed 10 cancer types with matched normal tissue data from GTEx (GTEx Consortium 2020)(Methods). Similar to the analysis of cancer tissues, we labeled each pair of chromosome-arm and CCL as recurrently gained (59 pairs), recurrently lost (45 pairs) or neutral (286 pairs), and updated the features associated with cancer types according to the CCL data (see Methods). We then applied the gain and loss ML models, which were trained on primary tumor data, to identify determinants of aneuploidy patterns of CCLs (Methods). The performance of the models was at least as good as for cancer types (gain model: auROC=83% and auPRC=49% (expected 15%); loss model: auROC=76% and auPRC=45% (expected 11%), Fig. 3A). These results indicate that similar factors affect aneuploidy in cancers and in CCLs, consistent with the highly similar aneuploidy patterns observed in tumors and in CCLs (Cohen-Sharir et al. 2021; Prasad et al. 2022).

**Figure 3.**
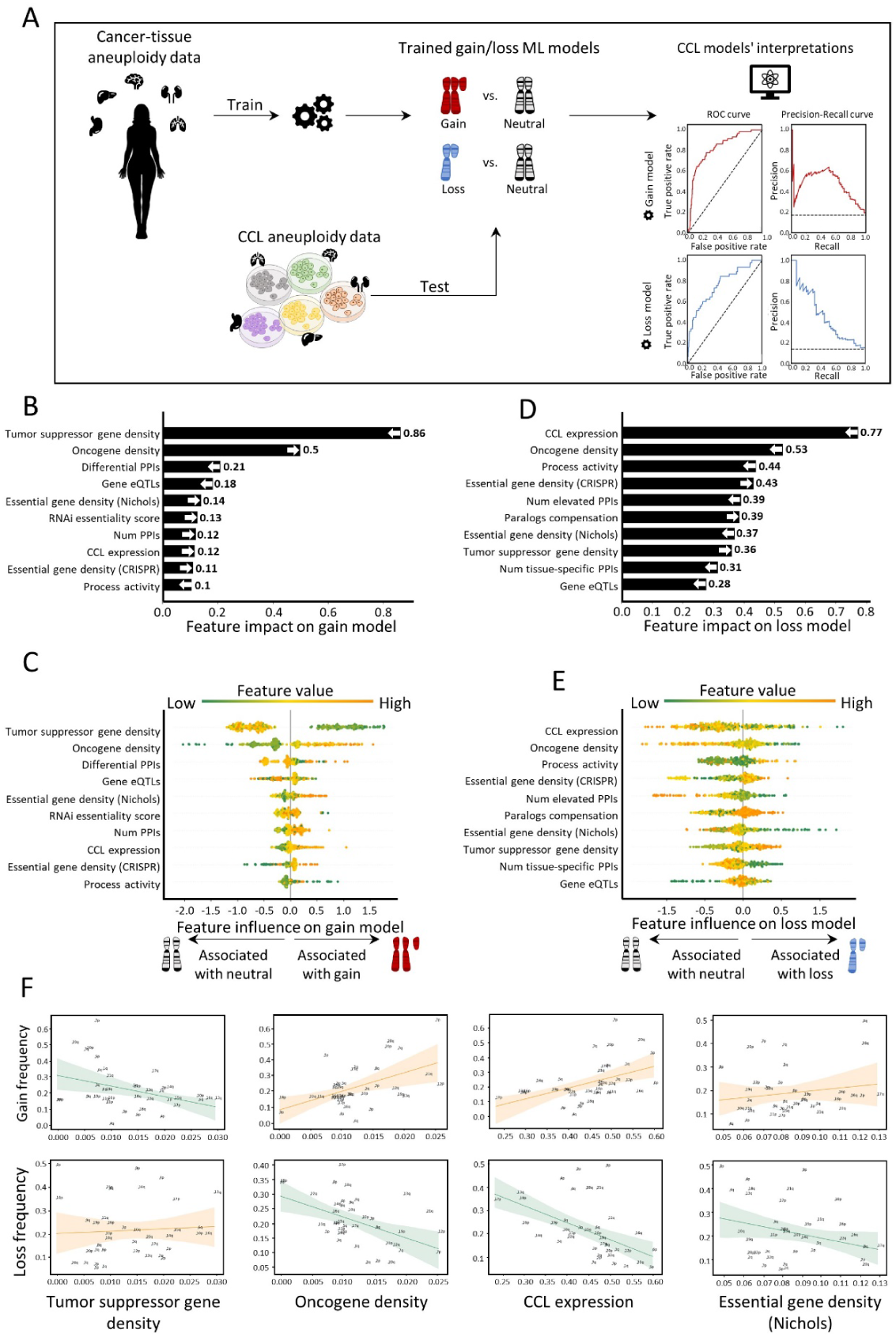
Aneuploidy patterns in CCLs and primary tumors are shaped by similar features. A. The ML scheme for analysis of aneuploidy patterns in CCL. The gain and loss models that were trained on aneuploidy patterns in primary tumors were applied to aneuploidy patterns in CCLs. Performance was measured using 10-fold cross-validation. Gain model (gradient boosting): auROC=83%, auPRC=49% (expected 15%). Loss model (XGBoost): auROC=76%, auPRC=45% (expected 11%). B. The average absolute contribution of the 10 topmost features to the gain model (see Fig. 2A). The order and directionality of the features generally agree with the gain model in primary tumors. C. A detailed view of the contribution of the 10 topmost features to the gain model (see Fig. 2B). D. Same as panel B for the loss model. The order and directionality of the features generally agree with the loss model in primary tumors. E. Same as panel C for the loss model. F. The correlation between top contributing features and the frequencies of chromosome-arms gain and loss, as measured by Spearman correlation. P-values were adjusted for multiple hypothesis testing using Benjamini-Hochberg procedure. Negative correlation between TSG density and gain frequency (ρ=-0.37, adjusted p=0.04). Positive correlation between TSG density and loss frequency (ρ=0.17, adjusted p=0.32). Positive correlation between OG density and gain frequency (ρ=0.44, adjusted p=0.012). Negative correlation between OG density and loss frequency (ρ=-0.28, adjusted p=0.13). Positive correlation between CCL expression and gain frequency (ρ=0.53, adjusted p=0.002). Negative correlation between CCL expression and loss frequency (ρ=-0.6, adjusted p=0.0006). Positive correlation between essential gene density and gain frequency (ρ=0.18, adjusted p=0.33). Negative correlation between essential gene density and loss frequency (ρ=-0.17, adjusted p=0.32).

We next used SHAP to assess the contribution of each feature to each of the models. ‘TSG density’ and ‘OG density’ remained the top contributing features for the gain model. Consistent with our results in primary tumors the contribution of ‘TSG density’ was much stronger than that of ‘OG density’, confirming the role of negative selection (Fig. 3B,C). In the loss model, the ranking of top features was slightly different than in primary tumors (Fig. 3D). Expression in CCL was the top feature, such that recurrently lost chromosome-arms were associated with lower gene expression in neutral cases. ‘OG density’ was one of the strongest contributing features for the loss model whereas ‘TSG density’ had weaker contribution, yet again in line with negative selection playing an important role in shaping cancer aneuploidy landscapes (Fig. 3D,E). Certain features of normal tissues were also highly ranked. The contribution of essential gene density was also consistent with the results in primary tumors (Fig. 3B,C).

As with the primary tumors, correlation analyses supported the contributions of the different features. CCL expression was highly correlated with chromosome-arm gain and anticorrelated with chromosome-arm loss (ρ=0.54, adjusted p=0.02, and ρ=-0.6, adjusted p=0.0006, respectively; Fig. 3F). Negative correlations were also observed between ‘TSG density’ and gain frequency (ρ=-0.37, adjusted p=0.04, Spearman correlation; Fig. 3F) and between ‘OG density’ and loss frequency (ρ=-0.28, adjusted p=0.1, Spearman correlation; Fig. 3F). Altogether, these results indicate that despite the continuous evolution of aneuploidy throughout CCL culture propagation (Ben-David et al. 2018), similar features drive aneuploidy recurrence patterns in primary tumors and in CCLs.

### Chromosome 13q aneuploidy pattern is governed by tissue-specific features, and KLF5 is a driver of 13q gain in colorectal cancer

In human cancer, a chromosome-arm is either recurrently gained or recurrently lost across cancer types, but is only rarely gained in some cancer types and lost in others (Ben-David et al. 2017; Taylor et al. 2018). An intriguing exception is chr13q. Of all chromosome-arms, chr13q is the chromosome-arm with the highest density of tumor suppressor genes (Fig. 2E). It is therefore not surprising that chr13q is recurrently lost across multiple cancer types (with a median of 30% of the tumors losing one copy of 13q across cancer types)(Ben-David et al. 2017; Taylor et al. 2018). Interestingly, however, chr13q is recurrently gained in human colorectal cancer (in 58% of the samples), suggesting that it can confer a selection advantage to colorectal cells in a tissue-specific manner. Indeed, when comparing colorectal cancer cell lines against cell lines of all other cancer types, chr13q was the top differentially-affected chromosome-arm (Fig. 4A,B). We therefore set out to study the basis for this unique tissue-specific aneuploidy pattern.

**Figure 4:**
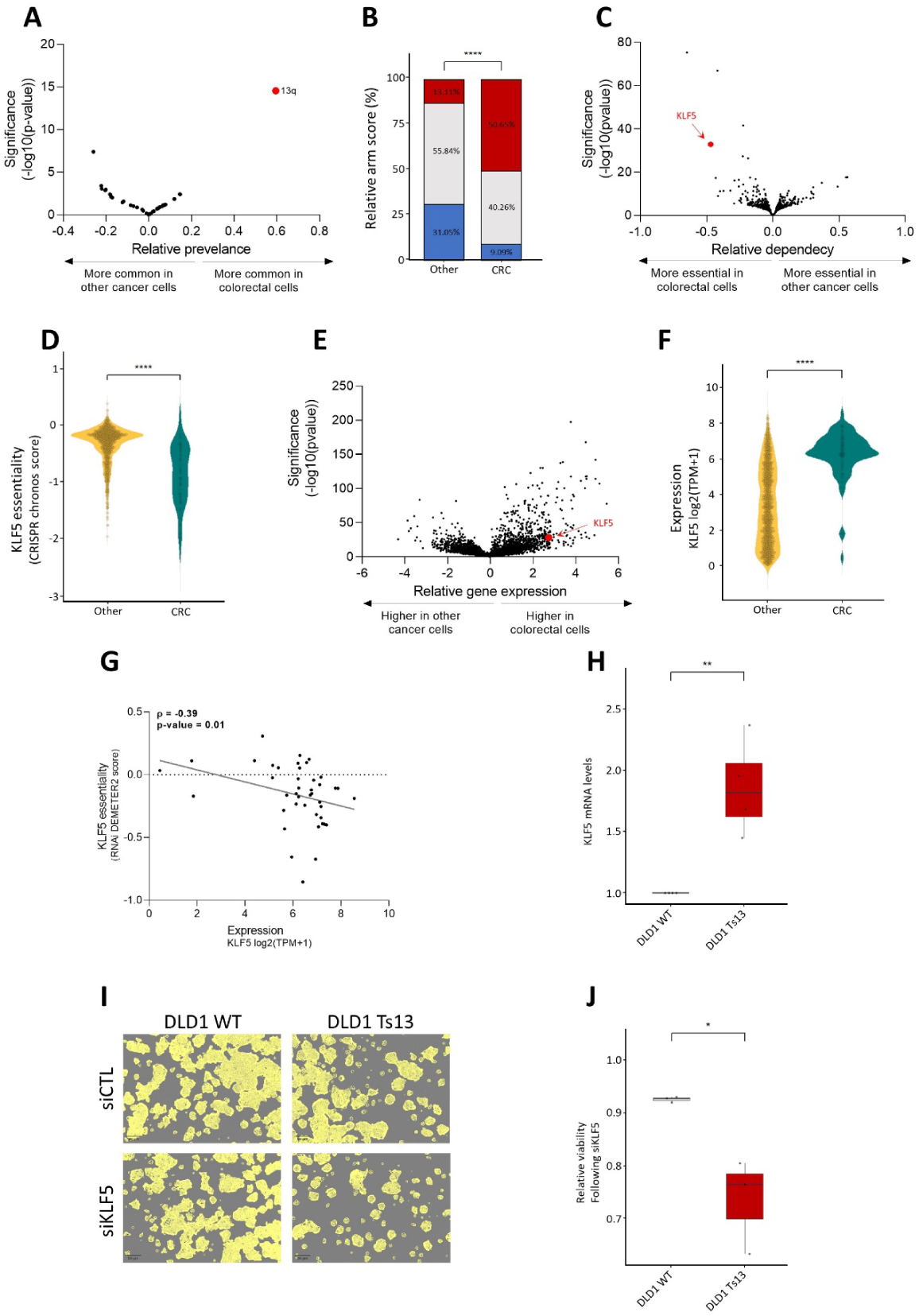
KLF5 is a potential driver of chromosome 13q gain in human colorectal cancer. A. Comparison of the prevalence of chromosome-arm aneuploidies in colorectal cancer cell lines against all other cancer cell lines. On the right side are the aneuploidies that are more common in colorectal cancer, and on the left side the ones that are less common in colorectal cancer. Chromosome-arm 13q (in red) is the top differential aneuploidy in colorectal cancer. B. Comparison of the prevalence of 13q aneuploidy prevalence between colorectal cancer cell lines and all other cancer cell lines. ****, p<0.0001; Chi-Square test. C. Genome-wide comparison of differentially essential genes between colorectal cancer cell lines and all other cancer cell lines. On the right side are the genes that are more essential in other cancer cell lines, an on the left side are those that are more essential in colorectal cancer, based on a genome-wide CRISPR/Cas9 knockout screen^1^. KLF5 (in red) is the second most differentially-essential gene in colorectal cancer cell lines. D. Comparison of the sensitivity to CRISPR knockout of KLF5 between colorectal cancer cell lines and all other cancer cell lines. ****, p<0.0001; two-tailed Mann-Whitney test. E. Genome-wide comparison of differentially expressed genes between colorectal cancer cell lines and all other cancer cell lines^1^. On the right side are the genes that are over-expressed in colorectal cancer and on the left side are those that are over-expressed in other cell lines. KLF5 (in red) significantly over-expressed in the colorectal cancer cell lines. F. Comparison of KLF5 mRNA levels between colorectal cancer cell lines and all other cancer cell lines. ****, p<0.0001; two-tailed Mann-Whitney test. G. Correlation between KLF5 mRNA expression and the sensitivity to KLF5 knockdown, showing that higher KLF5 expression is associated with increased sensitivity to its RNAi-mediated knockdown. ρ= -0.39, p=0.01; Spearman‘s correlation. H. Comparison of KLF5 mRNA levels between DLD1-WT (without trisomy of chromosome 13) and DLD1-Ts13 (with trisomy of chromosome 13) colorectal cancer cells. **, p= 0.0025; One-Sample t-test. I. Representative images of DLD1-WT and DLD1-Ts13 cells treated with siRNA against KLF5. DLD1-Ts13 cells were more sensitive to the knockdown. Cell masking (shown in yellow) was performed using live cell imaging (IncuCyte) for 72 hrs. Scale bar 100µm. J. Quantification of the relative response to KLF5 knockdown between DLD1-WT and DLD1-Ts13, as evaluated by quantifying cell viability in cells treated with siRNA against KLF5 vs. a control siRNA for 72 hrs. n=3 independent experiments. *, p=0.0346; one-sided paired t-test.

We performed a genome-wide comparison of differentially essential genes between colorectal cell lines and all other cell lines. The two top genes, which are much more essential in colorectal cancer cells than in other cancer types, were CTNNB1 and KLF5 (Fig. 4C). Of particular interest is KLF5, which is located on chr13q and colorectal cancer cell lines are significantly more sensitive to its knockout (Fig. 4D). KLF5 was reported to be tumor-suppressive in the context of several cancer types, such as breast and prostate (Chen et al. 2002; Ma et al. 2020). In colon cancer, however, not only is KLF5 important for tissue identity (Luo and Chen 2021), but it was also reported to be haploinsufficient (McConnell et al. 2009), potentially explaining why loss of chr13q is so rare in colorectal cancer. In line with a potential driving role in the recurrence of chr13q gain in colorectal cancer, KLF5 was among the most significantly overexpressed genes in colorectal cell lines vs. other cell lines (Fig. 4E,F) and its expression was significantly correlated with the sensitivity to its knockdown (Fig. 4G). To confirm the association between chr13q gain and KLF5 expression and dependency, we next turned to an isogenic system of human colon cancer cells (DLD1) into which trisomy 13 has been introduced (DLD1-T13) (Rutledge et al. 2016). Using this unique experimental system, we confirmed that trisomy 13 results in overexpression of KLF5 (Fig. 4H) and increased sensitivity to its siRNA-mediated genetic depletion (Fig. 4I,J and Fig. S7). We therefore propose that KLF5 contributes to the uniquely variable pattern of chr13q aneuploidy across cancer types.

### Paralog compensation is an important feature shaping tissue-specific aneuploidy patterns

One of the topmost contributing features to the loss model in primary tumors and in CCLs was ‘paralog compensation‘. This feature reflects whether the expression of a paralog is high relative to its homologous gene, suggesting that the paralog might compensate for the loss of that gene (Methods). Previous studies of hereditary disease genes showed that lower paralog compensation in a tissue was associated with disease manifestation (Barshir et al. 2018; Jubran et al. 2020). Paralog compensation was also shown in cancer tissues. In CCLs, essentiality of a gene was decreased with an increased expression of its paralog (Ito et al. 2021; Wang et al. 2015; Tsherniak et al. 2017). In primary tumors, paralog compensation was shown to be associated with increased prevalence of non-synonymous mutations (Zapata et al. 2018) and to correlate with the prevalence of homozygous gene deletion (Kegel and Ryan 2022). However, the contribution of paralog compensation to aneuploidy has not been studied to date.

Paralog compensation ranked 4^th^ and 6^th^ in the loss models of primary tumors and CCLs, respectively (Fig. 2C, Fig. 3D). In both, higher paralog compensation was associated with recurrently lost chromosome-arms, indicating that chromosome-arm loss can be tolerated better in tissues that highly express paralogs of this arm-residing genes (Fig. 2D, Fig. 3E). We also analyzed the correlations between paralog compensation and chromosome-arm loss and gain (Methods, Fig. 5A). Indeed, paralog compensation was positively correlated with the frequency of chromosome-arm loss in primary tumors (ρ=0.26, Spearman correlation) and in CCLs (ρ=0.46, adjusted p=0.01, Spearman correlation; Fig. S8A), and negatively correlated with the frequency of chromosome-arm gain in primary tumors (ρ=-0.21, Spearman correlation) and CCLs (ρ=-0.24, Spearman correlation; Fig. S8A).

**Figure 5:**
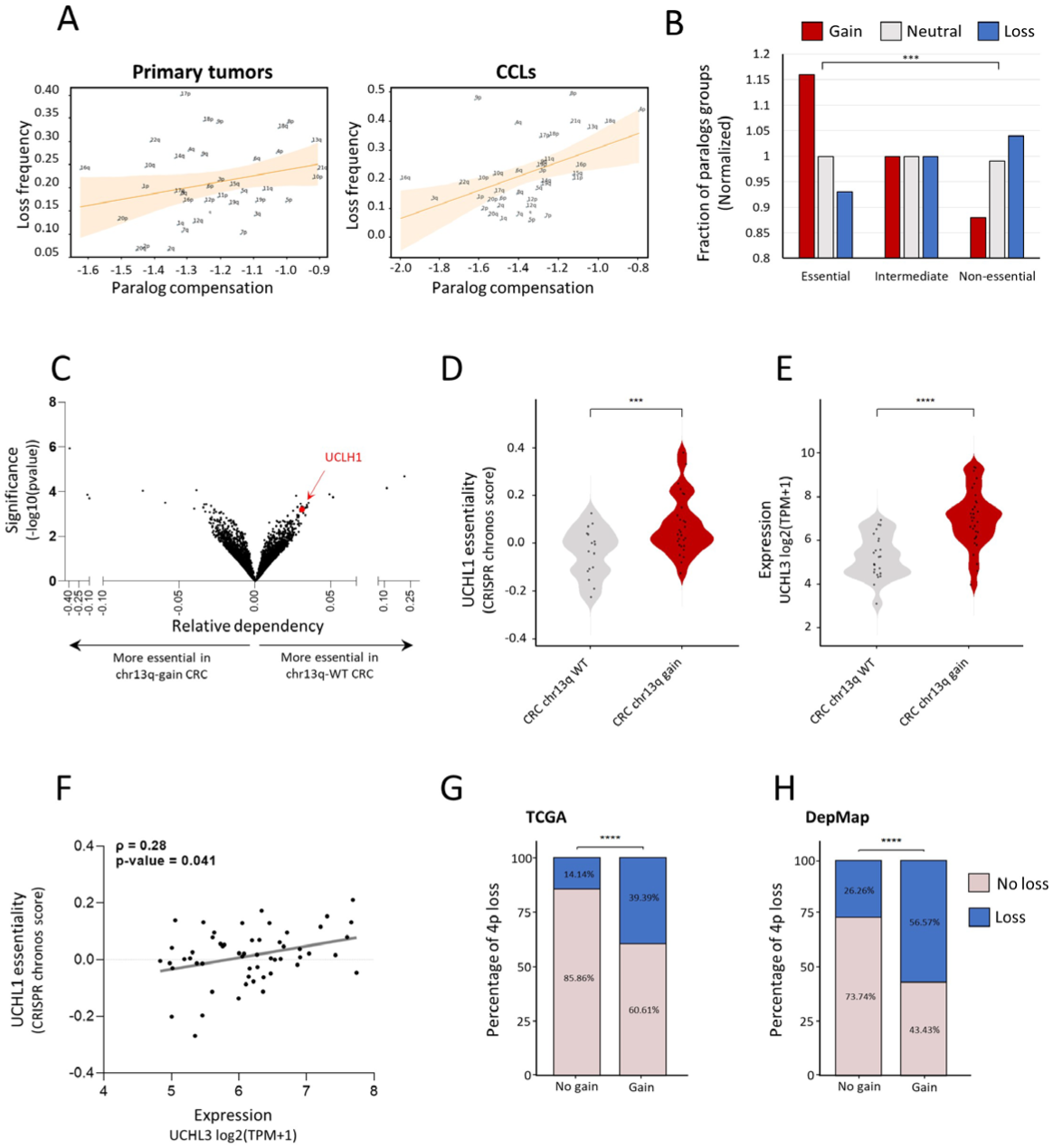
Paralog compensation is an important feature shaping tissue-specific aneuploidy patterns. A. The correlation between paralogs compensation values and loss frequency of chromosome arms in primary tumors (left, ρ = 0.26, adjusted p = 0.18, Spearman correlation) and in CCLs (right, ρ = 0.46, adjusted p = 0.01, Spearman correlation). B. A view into the aneuploidy patterns of paralogs of recurrently-lost genes. Recurrently-lost genes were divided into essential, intermediate, and non-essential genes. Paralogs of essential genes were more frequently gained, whereas paralogs of non-essential genes were more frequently lost. C. Genome-wide comparison of differentially essential genes in colorectal cell lines with chr13q gain vs. chr13q-WT colorectal cell line. On the right side are the genes that are more essential in chr13q-WT cells, and on the left side those that are more essential in chr13q-gain cells, based on a genome-wide CRISPR/Cas9 knockout screen^1^. UCHL1 (in red) is one of the top genes identified to be more essential in chr13q-WT cells. D. Comparison of the sensitivity to CRISPR knockout of UCHL1 between colorectal cell lines with and without chr13q gain. ***, p=0.0003; two-tailed Mann-Whitney test. E. Comparison of UCHL3 mRNA expression between colorectal cell lines with and without chr13q gain. ****, p<0.0001; two-tailed Mann-Whitney test. F. Correlation between UCHL3 mRNA expression and the sensitivity to UCHL1 knockout, showing that higher UCHL3 mRNA levels are associated with reduced sensitivity to UCHL1 knockout. ρ= 0.28, p=0.041; Spearman‘s correlation. G. Comparison of the prevalence of chr4p loss between human primary colorectal tumors with and without chr13q gain. ****, p<0.0001, Chi-square test. H. Comparison of the prevalence of chr4p loss between human colorectal cancer cell lines with and without chr13q gain. ****, p<0.0001, Chi-square test.

Next, we tested whether the effect of paralog compensation on the prevalence of chromosome-arm losses is associated with their essentiality. We therefore grouped recurrently lost genes into essential, intermediate, or non-essential genes, according to their essentiality in CCLs (Tsherniak et al. 2017) (Methods). We then assessed the prevalence of the chromosome-arm gain, loss or neutrality of their paralogs (Methods, Fig. S8B). The fraction of genes with neutral paralogs was similar in all essentiality groups (Fig. 5B). The fraction of genes with commonly gained paralogs was higher for essential genes, whereas non-essential genes were enriched for commonly lost paralogs (p=2.38e-24, Chi-squared test; Fig. 5B). The same trend was shown across various levels of essentiality (p=9.2e-16, KS test; Fig. S8C). Hence, paralog compensation, namely the gain or overexpression of the paralog, can facilitate chromosome-arm loss.

Next, we decided to identify a specific example. In human colon cancer, the long arm of chromosome 13 (chr13q) is commonly gained, as described above, whereas the short arm of chromosome 4 (chr4p) is commonly lost (Prasad et al. 2022; Taylor et al. 2018). We analyzed the association between chr13q-residing genes and the essentiality of their paralogs, revealing UCHL3 (chr13q)-UCHL1 (chr 4p) as the most significant correlation (Table S2 and Fig. 5C). Human colon cancer cell lines with chr13q gain were less sensitive to CRISPR/Cas9-mediated knockout of UCHL1 (Fig. 5D). Consistently, chr13q-gained cell lines had significantly higher mRNA levels of UCHL3 (Fig. 5E), and the expression of UCHL3 was significantly correlated with the essentiality of UCHL1 (Fig. 5F). We hypothesized that the relationship between these paralogs may affect the co-occurrence patterns of the chromosome-arms on which they reside. Indeed, both in primary human colon cancer and in colon cancer cell lines, loss of chr4p was significantly more prevalent when chr13q was gained (Fig. 5G,H). Together, these results demonstrate that paralog compensation can be affected by – and contribute to the shaping of – aneuploidy patterns.

## Discussion

Recurrent aneuploidy patterns are an intriguing phenomenon that is only partly understood. Previous efforts to explain this phenomenon focused on specific aspects, such as OG and TSG densities (Sack et al. 2018; Taylor et al. 2018) or gene expression levels (Patkar et al. 2021), which were interrogated using statistical methods and correlation analyses. Here, in contrast, we approached this phenomenon using ML. As with other ML applications, it allowed us to study multiple aspects simultaneously. Yet, unlike classical ML-based studies that mainly aim to improve prediction, for example by using deep learning to predict gene dependency in tumors (Chiu et al. 2021), our focus was on interpretability. In fact, we built chromosome-arm gains and loss models only to then identify factors that shape aneuploidy patterns. Interpretable ML was recently applied to reveal genetic attributes that contribute to the manifestation of Mendelian diseases (Simonovsky et al. 2023). In this study we applied it for the first time in the context of aneuploidy, or at chromosome-arm resolution.

The capability of ML to concurrently assess multiple features opened the door for assessing the relevance of features that have not been rigorously studied to date, such as paralogs compensation. Yet, ML has its limitations. Mainly, the number of features that could be analyzed depends on the size of the labeled dataset (Hua et al. 2005), which, in aneuploidy, was restricted by the numbers of chromosome-arms and cancer types. We therefore analyzed 20 types of features, and tested linear regression and tree-based ML methods, which, unlike deep learning, are suitable for this size of data. Following prediction, our main goal was to assess the relative contribution of each feature to the model‘s decision and its directionality using SHAP. However, SHAP results should be interpreted with caution. Due to the hierarchical nature of decision trees, features that are located low in the decision tree explain only a small fraction of the cases. To estimate feature directionality more broadly, we then explicitly correlated between feature values and chromosome-arm gain and loss frequency.

The features that we studied included known and previously underexplored attributes of chromosome-arms and of healthy and cancer tissues and cells (Fig. 1A,B). OG and TSG densities, which have previously been observed to be enriched on gained and lost chromosome-arms, respectively (Davoli et al. 2013; Sack et al. 2018), were top contributing features in both models, thereby supporting the validity of our approach (Fig 2A,C). In the gain model in particular, their contribution was over 2.6 and 5 times stronger, respectively, than any other feature (Fig. 2A). As our TSG and OG features were cancer-independent, their importance may explain the observation that certain chromosome-arms tend to be either gained or lost across multiple cancer types (Ben-David et al. 2017; Taylor et al. 2018). Their relative contribution, however, was surprising. In both models, negative associations were much stronger than positive associations: OG density contributed to chromosome-arm loss more than TSG density, implying that it was more important to maintain OGs than to lose TSG (Fig. 2B,D). The reciprocal relationship was true for chromosome-arm gain, as it was more important to maintain TSGs than to gain OGs (Fig. 2A,C). These results were validated using correlation analyses (Fig. 2E), and were recapitulated in CCLs (Fig. 3). They highlight the importance of negative selection for shaping cancer aneuploidy landscapes (Shih et al. 2023; Ben-David and Amon 2020).

A known factor that contributed to both models was gene expression in primary tumors (‘TCGA expression’, Fig. 2) and in CCLs (‘CCL expression’, Fig. 3). This result suggests that cancers tend to gain chromosome-arms that are enriched for highly-expressed genes, and tend to lose chromosome-arms that are enriched for lowly-expressed genes. A Similar trend was shown recently for gene expression in normal tissues (Patkar et al. 2021). Our approach was capable of comparing the relative contributions of both features. We found that the contribution of gene expression in normal tissue was lower than in cancer tissues, as also evident in its lower correlation with the frequencies of chromosome-arm gain and loss (Fig. S5). Nevertheless, other features that were derived from gene expression in normal tissues ranked highly, such as the number of PPIs in the gain model and paralog compensation in the loss model, and hence expression in normal tissues is also important (Fig. 2).

A previously under-explored feature that we considered was paralog compensation. Paralog compensation was shown to play a role in the manifestation of Mendelian and complex diseases (Barshir et al. 2018; Jubran et al. 2020) and in the dispensability of genes in tumors (Kegel and Ryan 2022; Zapata et al. 2018) and CCLs (Ito et al. 2021; Tsherniak et al. 2017; Wang et al. 2015), but was not studied in the context of aneuploidy. Here, paralog compensation was among the top contributors to the loss model (Fig. 2C, Fig. 3D). The directionality of this feature and correlation analyses showed that, relative to genes located on neutral chromosome-arms, genes located on lost chromosome-arms tend to have higher compensation by paralogs (Fig. 5A). This suggests that chromosome-arm loss is facilitated, or better tolerated, through paralogs’ expression. We also showed that the more essential recurrently lost genes are, the more likely they are to be associated with gains of paralog-bearing chromosome-arms (Fig. 5B). We further demonstrated this for a specific example (the UCHL3-UCHL1 paralog pair; Fig. 5). Overall, our analysis reveals that compensation between paralogs though expression or chromosome-arm gain plays an important role in shaping the landscape of chromosome-arm loss.

Combining the different results, our models reveal a previously under-appreciated role for negative selection in driving human cancer aneuploidy. This was evident by the tendency not to lose chromosome arms with high OG density, high frequency of essential genes, or low compensation by paralogs, and not to gain chromosome arms with high TSG density (Fig. 6). Previous studies have shown that positive selection outweighs negative selection in shaping the point mutation landscape of human tumors (Martincorena et al. 2017). However, the strong fitness cost associated with aneuploidy suggests that the aneuploidy landscape of tumors might be strongly affected by negative selection as well (reviewed in (Ben-David and Amon 2020)). Interestingly, evidence for the involvement of negative selection in shaping the copy number alteration (CNA) landscapes of tumors has been proposed in a recent preprint that analyzed CNA length distributions across human tumors (Shih et al. 2023). Our study lends further independent support to the importance of negative selection in shaping the landscape of aneuploidy across human cancers (Fig. 6).

**Figure 6:**
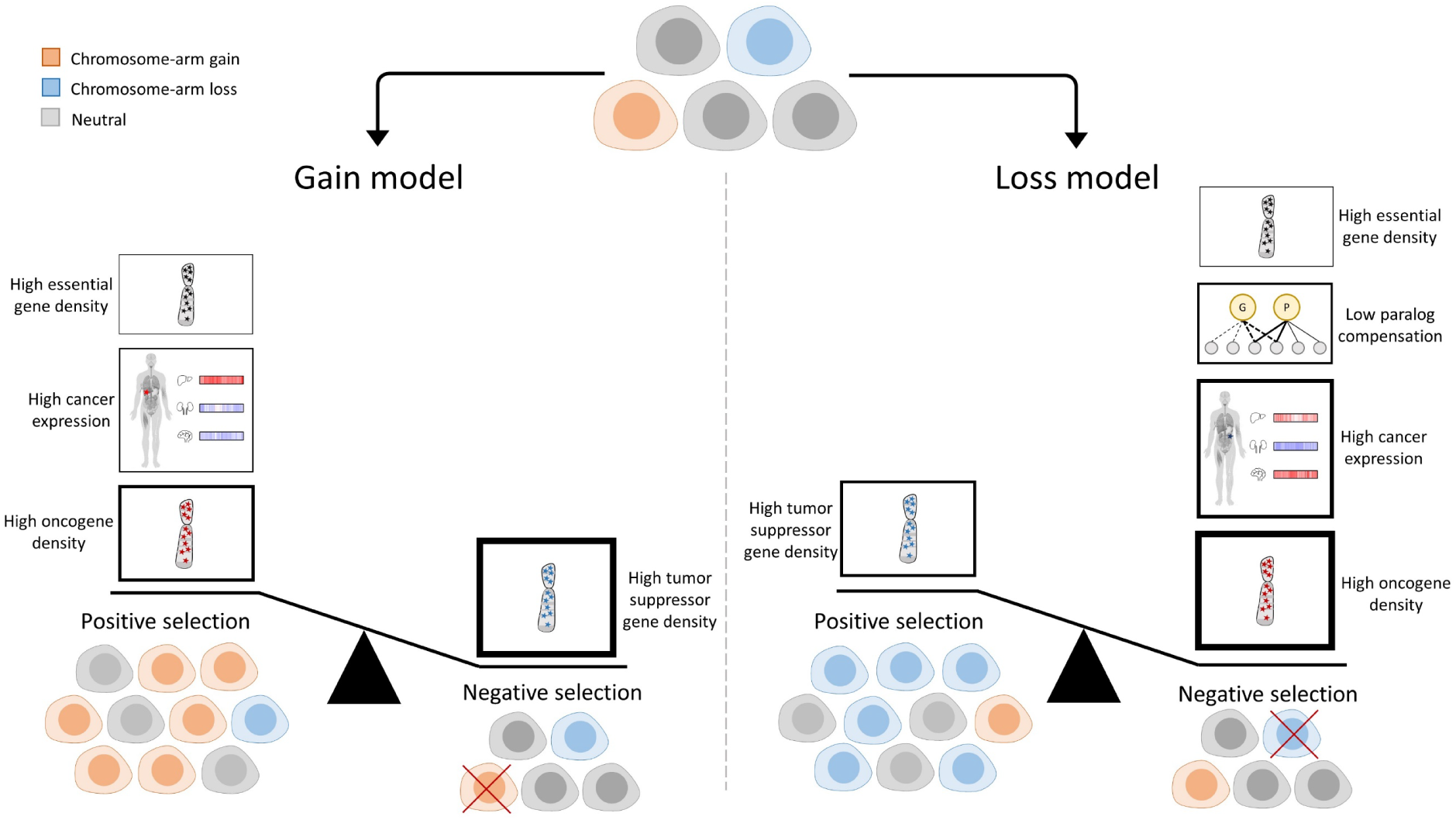
A schematic presentation of the study. Cancer evolution is shaped by negative and positive selection leading to enrichment or depletion of cells with distinct aneuploidy patterns. In the gain model (left), main contributors to positive selection of gained chromosome arms are: (1) high oncogene density; (2) high expression of genes in the cancer tissue; and (3) high essential gene density. A major contributor to negative selection is high tumor-suppressor gene density. Importantly, the density of TSGs is more important than the density of OGs for predicting chromosome-arm gains. In the loss model (right), a main contributor to positive selection of lost chromosome arms is high tumor-suppressor gene density. Major contributors to negative selection are high oncogene density, high expression of genes in the cancer tissue, low compensation by paralogs, and high density of essential genes. In both models, the features associated with negative selection have higher overall contribution than features associated with positive selection. The thickness of the borders of the boxes reflect the relative contribution of the features to the model.

Lastly, we explored one example of a unique aneuploidy pattern (chr13q) that is recurrently altered in opposite directions in different cancer types. In line with tumor suppressors and oncogenes being a major feature explaining aneuploidy patterns, we identified KLF5 as a colorectal-specific dependency gene. Using an isogenic system of colorectal cancer cells with/without gain of chr13, we experimentally demonstrated that this aneuploidy is associated with increased expression and increased essentiality of KLF5. We therefore propose that KLF5 might explain why chr13q is commonly gained and rarely lost in colorectal cancer, unlike its recurrent loss across multiple other cancer types.

Overall, our study provides novel insights into the forces that shape the tissue-specific patterns of aneuploidy observed in human cancer, and demonstrates the value of applying ML approaches to dissect this complicated question. Our results suggest that aneuploidy patterns are shaped by a combination of tissue-specific and non-tissue-specific factors. Negative selection in general, and paralog compensation in particular, play a major role in shaping the aneuploidy landscapes of human cancer, and should therefore be computationally modeled and experimentally studied in the research of cancer aneuploidy.

## Methods

### Chromosome-arm aneuploidy patterns per cancer

Aneuploidy patterns were available for all (39) chromosome-arms in 33 cancer types from Genome Identification of Significant Targets in Cancer 2 (GISTIC2) (Mermel et al. 2011). A chromosome-arm was considered as gained or lost in a specific cancer if the q-value of its amplification or deletion, respectively, was smaller than 0.05 (in case of ties, decision was made based on the lower q-value). Otherwise, the chromosome-arm was considered as neutral.

### Construction of a features dataset of chromosome-arm and cancer type pairs

For each chromosome-arm and cancer, we created features that were inferred from data of chromosome-arms, genes, cancer tissues and CCLs, and normal tissues (Fig. 1B, Table S1). A schematic pipeline of the dataset construction appears in Fig. S1. The different types of features are described below.

#### Features of chromosome-arms

Each chromosome-arm was associated with three types of features, including oncogene density, tumor suppressor gene density, and essential gene density. Oncogene density and tumor suppressor gene density per chromosome-arm were obtained from davoli et al. (Davoli et al. 2013). Data of essential genes was obtained from Nichols et al (Nichols et al. 2020), where a gene was considered essential if its essentiality probability was > 0.8. The density of essential genes per chromosome-arm was calculated as the fraction of essential genes out of the protein-coding genes on that chromosome-arm. Next, we associated each pair of chromosome-arm and cancer with features of that chromosome-arm.

#### Features of cancer tissues

Each pair of chromosome-arm and cancer type was associated with four types of cancer-related features, including transcriptomics, essentiality by CRISPR or RNAi in CCLs, and cancer-specific density of essential genes. Transcriptomics was based on transcriptomic profiles of 33 cancer types from TCGA (Xena browser, (Goldman et al. 2020)). Per cancer, we associated each gene with its median expression level in samples of that cancer. To avoid expression bias due to chromosome-arm gain or loss, the median expression of each gene was computed from samples where the chromosome-arm harboring the gene was neutral (Taylor et al. study (Taylor et al. 2018). Essentiality by CRISPR was based on CRISPR screens of 24 CCLs from DepMap (Tsherniak et al. 2017). Essentiality by RNAi was based on RNAi data of 22 CCLs from DepMap (Tsherniak et al. 2017) In each of these datasets, the score of each gene indicated the change, relative to control, in the growth rate of the cell line upon gene inactivation via CRISPR or RNAi. Accordingly, genes with negative scores were essential for the growth of the respective cell line. We associated each gene with its median essentiality score based on either CRISPR or RNAi per cell line. To reflect gene essentiality more intuitively, we reversed the direction of the scores (multiplied them by -1), so that more essential genes had higher scores. To avoid bias due to chromosome-arm gain or loss, the median essentially of each gene was computed from samples where the chromosome-arm harboring the gene was neutral (Taylor et al. 2018).Cancer-specific density of essential genes was calculated as the fraction of essential genes (CRISPR-based essentiality score > 0.5) in a given CCL out of the protein-coding genes residing on that chromosome-arm.

#### Features of normal tissues

Each pair of chromosome-arm and cancer type was associated with 13 types of features that were derived from (Simonovsky et al. 2023). We associated each cancer type with the normal tissue in which it originates (Table S1).

##### Transcriptomics

Data of normal tissues included transcriptomic profiles of 54 adult human tissues measured via RNA-sequencing from GTEx v8 (GTEx Consortium 2020). Each gene was associated with its median expression in each adult human tissue. Genes with median TPM > 1 in a tissue were considered as expressed in that tissue.

##### Tissue-specific genes

Per gene, we measured its expression in a given tissue relative to other tissues using z-score calculation. Genes with z-score > 2 were considered tissue-specific. Lastly, we associated each chromosome-arm and tissue with the density of tissue-specific genes.

##### PPI features

Each gene was associated with the set of its PPI partners. We included only partners with experimentally-detected interactions that were obtained from MyProteinNet web-tool (Basha et al. 2015). Per each tissue we associated each gene with four PPI-related features: (i) ‘Number PPIs’ was set to the number of PPI partners that were expressed in that tissue. (ii) ‘Number elevated PPIs’ relied on preferential expression scores computed according to (Basha et al. 2020), and was set to the number of PPI partners that were preferentially expressed in that tissue (preferential expression > 2, (Sonawane et al. 2017). (iii) ‘Number tissue-specific PPIs’ was set to the number of PPI partners that were expressed in that tissue and in at most 20% of the tissues. (iv) ‘Differential PPIs’ relied on differential PPI scores per tissue from The DifferentialNet Database (Basha et al. 2020), and was set to gene‘s median differential PPI score per tissue. If the gene was not expressed in a given tissue, its feature values in that tissue were set to 0.

##### Differential process activity features

Differential process activity scores per gene and tissue were obtained from (Sharon et al. 2022). The score of a gene in a given tissue was set to the median differential activity of the Gene Ontology (GO) processes involving that gene. The differential activity was relative to the activity of the same processes in other tissues.

##### eQTL features

eQTLs per gene and tissue were obtained from GTEx (GTEx Consortium 2020). Each gene was associated with the p-value its eGene in that tissue.

##### Paralog compensation features

Each gene was associated with its best matching paralog according to Ensembl-BioMart. Per tissue, the gene score was set to the median expression ratio of the gene and its paralog, as described in (Barshir et al. 2018; Jubran et al. 2020). Accordingly, high values mark genes with low paralog compensation.

##### Development features

Transcriptomic data of seven human organs measured at several time points during development were obtained from (Cardoso-Moreira et al. 2019). We united time points into time periods including fetal (4-20 weeks post-conception), childhood (newborn, infant, and toddler) and young (school, teenager and young adult). Per organ, we associated each gene with its median expression level per period. Next, we created an additional feature that reflected the expression variability of each gene across periods.

#### Transforming gene features into chromosome-arm features

Some of the features described above referred to genes. To create chromosome-arm-based features, we grouped together genes that were located on the same chromosome-arm (Cunningham et al. 2022). Next, to highlight differences between tissues, for each feature, we associated a gene with its value in that tissue relative to other tissues. Features that were already tissue-relative, including ‘Differential PPIs’ and ‘Differential process activity’, were maintained. Other features were converted into tissue-relative values via a z-score calculation (see Equation 1). Lastly, per feature, we ranked genes by their tissue-relative score, and associated each chromosome-arm with the median score of the genes ranking at the top 10% (Fig. S2). Transcriptomic features in testis and whole blood were highly distinct from other tissues, we normalized all transcriptomic features per tissue. To reflect paralog compensation more intuitively, we reversed the direction of the resulting features (multiplied them by -1), so that genes with higher compensation had higher scores.

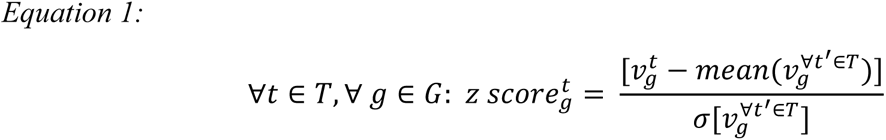

*T* denotes the set of tissues; *G* denotes the set of genes; *v* denotes the value of the feature; ᓂ denotes the standard deviation.

#### Construction of the final dataset

The features described above referred to chromosome-arms in cancers, CCLs, and normal tissues. To create chromosome-arm features per cancer, we associated each cancer with the chromosome-arm features of its tissue of origin and CCL (Table S1). In case several tissues were associated with the same cancer (e.g., skin sun-exposed and not sun-exposed), we set the cancer and chromosome-arm value to their median chromosome-arm values. The final dataset contained features for all pairs of 39 chromosome-arms and 24 cancers for which the tumor‘s normal tissue of origin was available in GTEx (Table S1).

### ML application to model chromosome-arm and cancer aneuploidy

Below we describe the ML method used for aneuploidy classification and the SHAP (SHapley Additive exPlanations) analysis of feature importance that was used to interpret the resulting models.

#### Aneuploidy ML classification models

We constructed two ML models: a gain model that compared between gained and unchanged (neutral) chromosome-arms; and a loss model that compared between lost and unchanged (neutral) chromosome-arm.

#### ML comparison and implementation

Per model, we tested several ML methods, including logistic regression, XGBoost, gradient boosting, random forest and bagging. All ML methods were implemented using the Scikit-learn python package (Pedregosa et al. 2011), except for XGB, which was implemented using the Scikit-learn API of the XGBoost package (Chen and Guestrin 2016) . To assess the performance of each model, we used 10-fold cross validation. Then, we calculated the au-ROC and the au-PRC. Each point on the curve corresponded to a particular cutoff that represented a trade-off between sensitivity and specificity and between precision and recall, respectively.

#### SHAP analysis of feature importance

To measure the importance and contribution of the different features we used SHAP (Shapley Additive exPlanations) algorithm (Lundberg et al. 2020). Per model, we created a SHAP summary and a SHAP feature importance plot. To visualize the direction of each feature (i.e., whether high values correspond to gain/loss or opposingly, to neutral), we considered the top 50% values of a given feature and calculated their average SHAP values. Similar results were obtained upon modeling chromosome-arm gain versus no-gain (i.e., neutral and lost chromosome-arms), and chromosome-arm loss versus no-loss (i.e., neutral and gained chromosome-arms), and repeating the procedures described above (Fig. S6).

### Correlation analysis

We correlated between feature values and the frequency of chromosome-arm gain or loss. The frequency of chromosome-arm gain/loss in cancers was obtained from GISTIC2 (Mermel et al. 2011). The frequency of chromosome-arm gain/loss in CCLs were obtained from (Prasad et al. 2022). Per chromosome-arm, its gain (loss) frequency was set to the median gain (loss) across cancers or CCLs. The feature value was set to median across cancers or CCLs. We used Spearman correlation, and p-values were adjusted using Benjamini-Hochberg procedure (Benjamini and Hochberg 1995).

### Paralog compensation analysis

For each cancer type and chromosome-arm, we considered all paralog pairs in which one of the genes resides on that chromosome-arm. We focused on recurrently-lost genes per cancer type as defined by GISTIC2 (Mermel et al. 2011). We divided those genes by their minimal CRISPR essentiality score in CCLs that match the same cancer type (Table S1). Genes with a score ≤-0.5 were considered essential, and genes with a score ≥-0.3 were considered non-essential. Other genes were considered intermediate. Per gene, we checked whether its paralog was recurrently gained, lost, or neutral, in the same cancer, as detailed in Fig. S8B.

### Chromosome-arm aneuploidy patterns in CCLs

Aneuploidy patterns were available for all (39) chromosome-arms in 14 CCLs from (Prasad et al. 2022). A chromosome-arm was considered as gained or lost in a CCL if the q-value of its amplification or deletion, respectively, was smaller than 0.15 (in case of ties, decision was made based on the lower q-value). Otherwise, the chromosome-arm was considered as neutral. *Construction of a feature dataset of chromosome-arm and CCL pairs:* The features dataset was similar to the dataset created for cancers, with the following exceptions. In features of cancer tissues, we replaced the transcriptomic features of cancers with transcriptomic features of CCLs. We obtained transcriptomic data of 25 CCLs from DepMap (Tsherniak et al. 2017), and constructed the feature values per chromosome-arm and CCL as described above per chromosome-arm and cancer. Development features were removed since only a small number of CCLs had a matching organ. The final dataset contained features for all pairs of 39 chromosome-arms and 10 CCLs for which the tumor‘s normal tissue of origin was available in GTEx.

### Cell culture

DLD1-WT cells and DLD1-Ts13 cells were cultured in RPMI-1640 (Life Technologies) with 10% fetal bovine serum (Sigma-Aldrich) and 1% penicillin-streptomycin-glutamine (Life Technologies). Cells were incubated at 37 °C with 5% CO2 and passaged twice a week using Trypsin-EDTA (0.25%) (Life Technologies). Cells were tested for mycoplasma contamination using the MycoAlert Mycoplasma Detection Kit (Lonza), according to the manufacturer‘s instructions.

### qRT-PCR

Cells were harvested using Bio-TRI® (Bio-Lab) and RNA was extracted following manufacturer‘s protocol. cDNA was amplified using GoScript™ Reverse Transcription System (Promega) following manufacturer‘s protocol. qRT-PCR was performed using Sybr® green, and quantification was performed using the ΔCT method. The following primer sequences were used: human KLF5, forward, 5’ ACACCAGACCGCAGCTCCA 3’ and reverse 5’ TCCATTGCTGCTGTCTGATTTGTAG 3‘.

### siRNA transfection

For siRNA experiments, cells were plated in 96-well plates at 6,000 cells per well and treated with compounds 24 hrs later. The cells were transfected with 15nM siRNA against KLF5 (ONTARGETplus SMART-POOL®, Dharmacon), or with a control siRNA at the respective concentration (ONTARGETplus SMART-POOL®, Dharmacon) using Lipofectamine® RNAiMAX (Invitrogen) following the manufacturer‘s protocol. Cell proliferation following siRNA transfection was followed by live cell imaging using Incucyte® (Satorius). The effect of KLF5 knockdown on cell viability/proliferation was measured by the MTT assay (Sigma M2128) at 72hrs (or at the indicated time point) post-transfection. 500ug/mL MTT salt was diluted in complete medium and incubated at 37°C for 2 hrs. Formazan crystals were extracted using 10% Triton X-100 and 0.1N HCl in isopropanol, and color absorption was quantified at 570nm and 630nm (Alliance Q9, Uvitec).

### Cancer cell line data analysis

mRNA gene expression values, arm-level CNAs, CRISPR and RNAi dependency scores (Chronos and DEMETER2 scores, respectively) were obtained from DepMap 22Q4 release (www.depmap.org). Effect size, p-values and q-values (Fig. 4A,C,E, Fig. 5C) were taken directly from DepMap and were calculated as described in Tsherniak et al. Effect size, Spearman‘s R and p-values in Fig. 4G and Fig. 5F were calculated using R functions.

The analyses that led to our choice of the paralog pair UCHL3-UCHL1 are summarized in Table S2. In the left column are the paralogs that reside on chr-13q, which is frequently gained; in the adjacent column are the respective paralogs that reside on commonly lost chromosomes. The following columns describe the Spearman correlation between each paralog pair and the respective p-value. The right-hand columns describe the effect size of chr-13q paralogs’ gene expression between CRC cell lines with and without chr13q gain. Our criteria for finding appropriate paralog pairs were as follows: firstly, to have a high expression of the chr-13q paralogs in CRC cell lines. Secondly, we aimed to reach a significant correlation between chr13q-residing genes and the essentiality of their paralogs.

### Statistical analyses

Statistical analysis was performed using GraphPad PRISM® 9.1 software. Details of the statistical tests were reported in figure legends. Error bars represent SD. All experiments were performed in at least three biological replicates.

## Acknowledgments

The authors would like to thank Jason Sheltzer for providing DLD1-WT and DLD1 Ts13 cell lines.

J.J. wishes to thank the Baroness Ariane de Rothchild Women Doctoral Program.

## Supplementary figures

**Figure S1.**
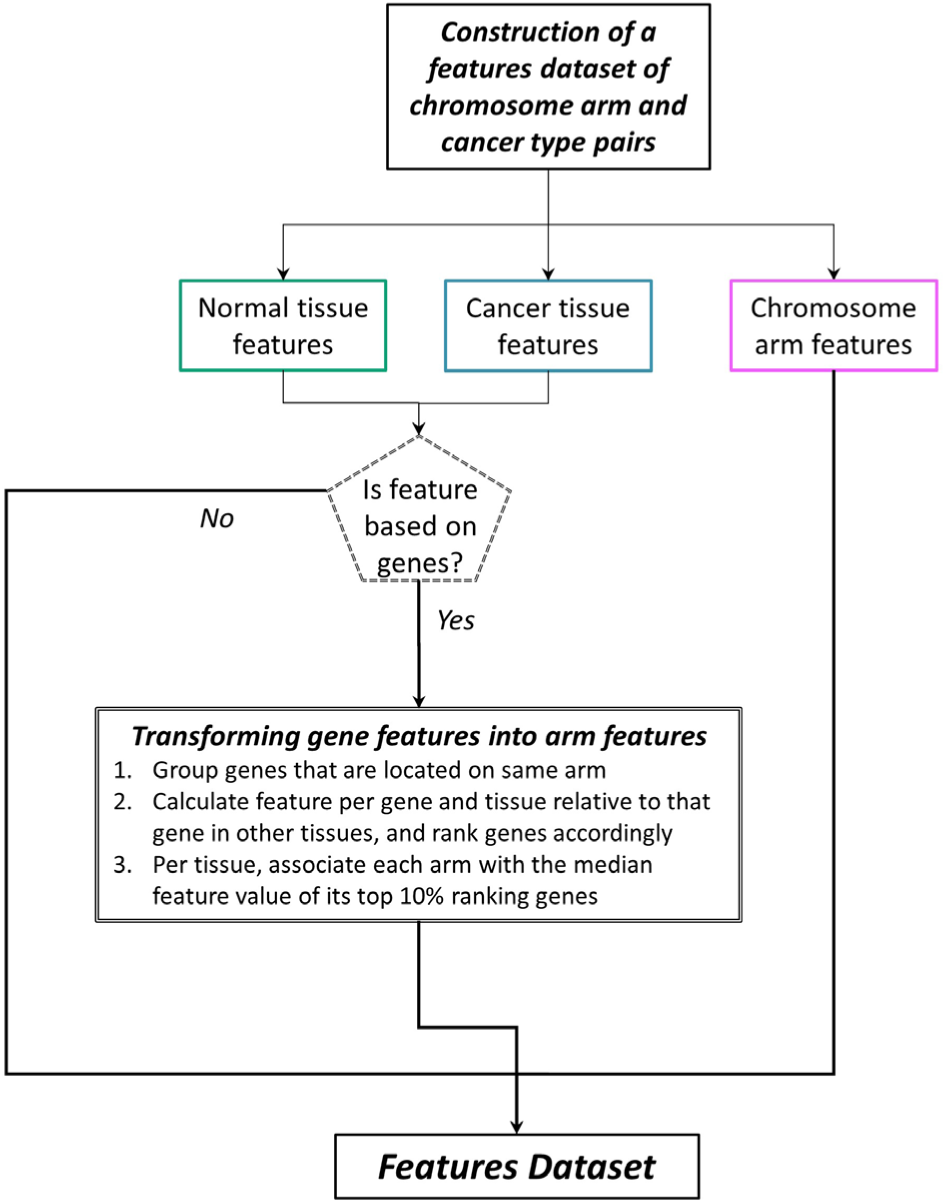
Workflow for constructing the features dataset. We associated each pair of chromosome-arm and cancer type with three categories of features: chromosome-arm, normal tissues and cancer tissues features. Features belonging to the chromosome-arm category were independent of cancer types. Features belonging to the categories of normal and cancer tissue features were either inferred from data of the entire chromosome-arm, in which case no transformation was needed, or were inferred from data of individual genes, in which case these data were collated per chromosome-arm and transformed into chromosome-arm-based features.

**Figure S2.**
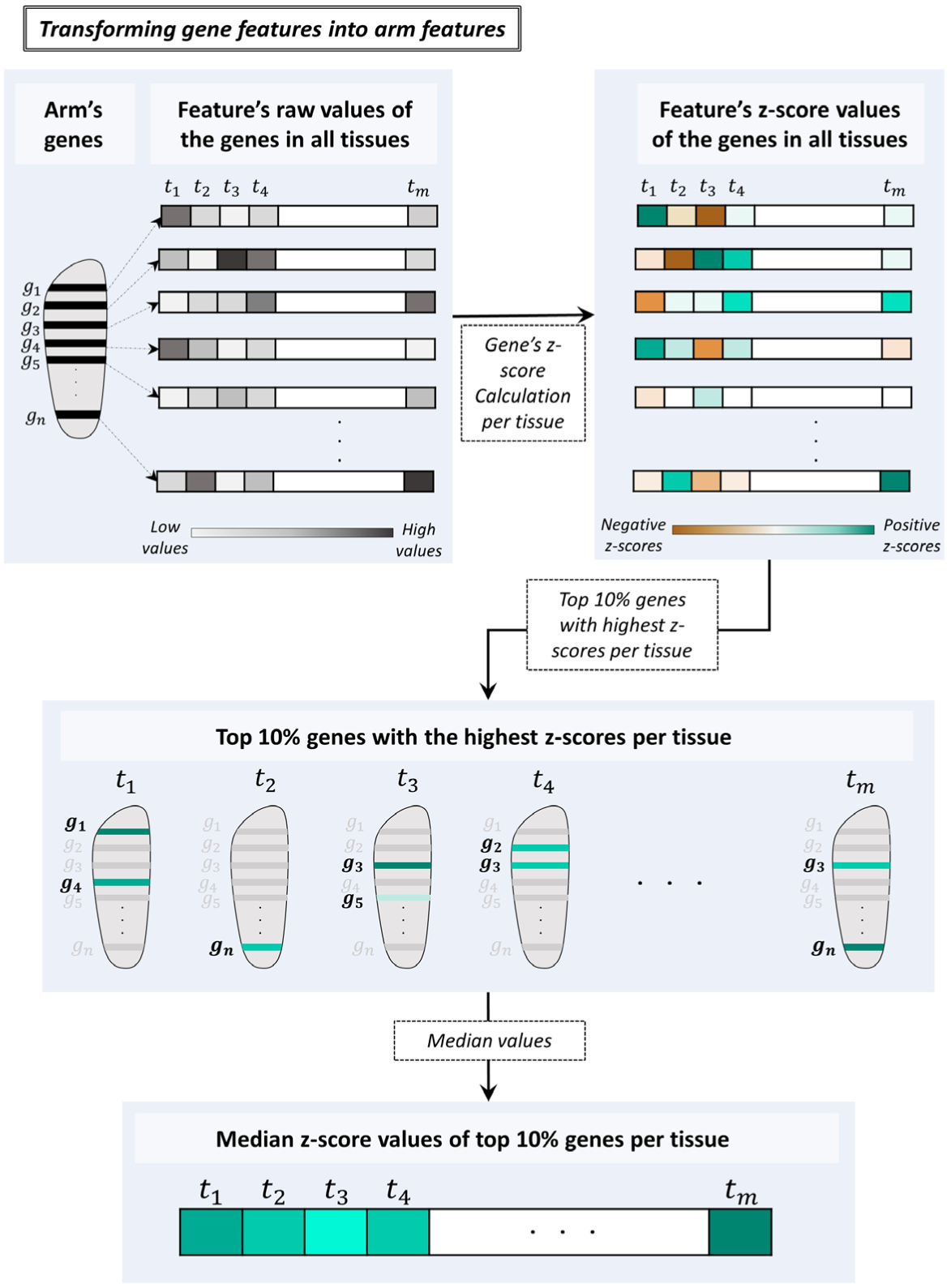
Workflow for transforming gene-related features into chromosome-arm related features. For each gene-related feature and chromosome-arm, genes located on that chromosome-arm were grouped. Their raw values in the different tissues were transformed into z-scores per gene and tissue. Then, each tissue was associated with the median of the top 10% genes with the highest z-score.

**Figure S3.**
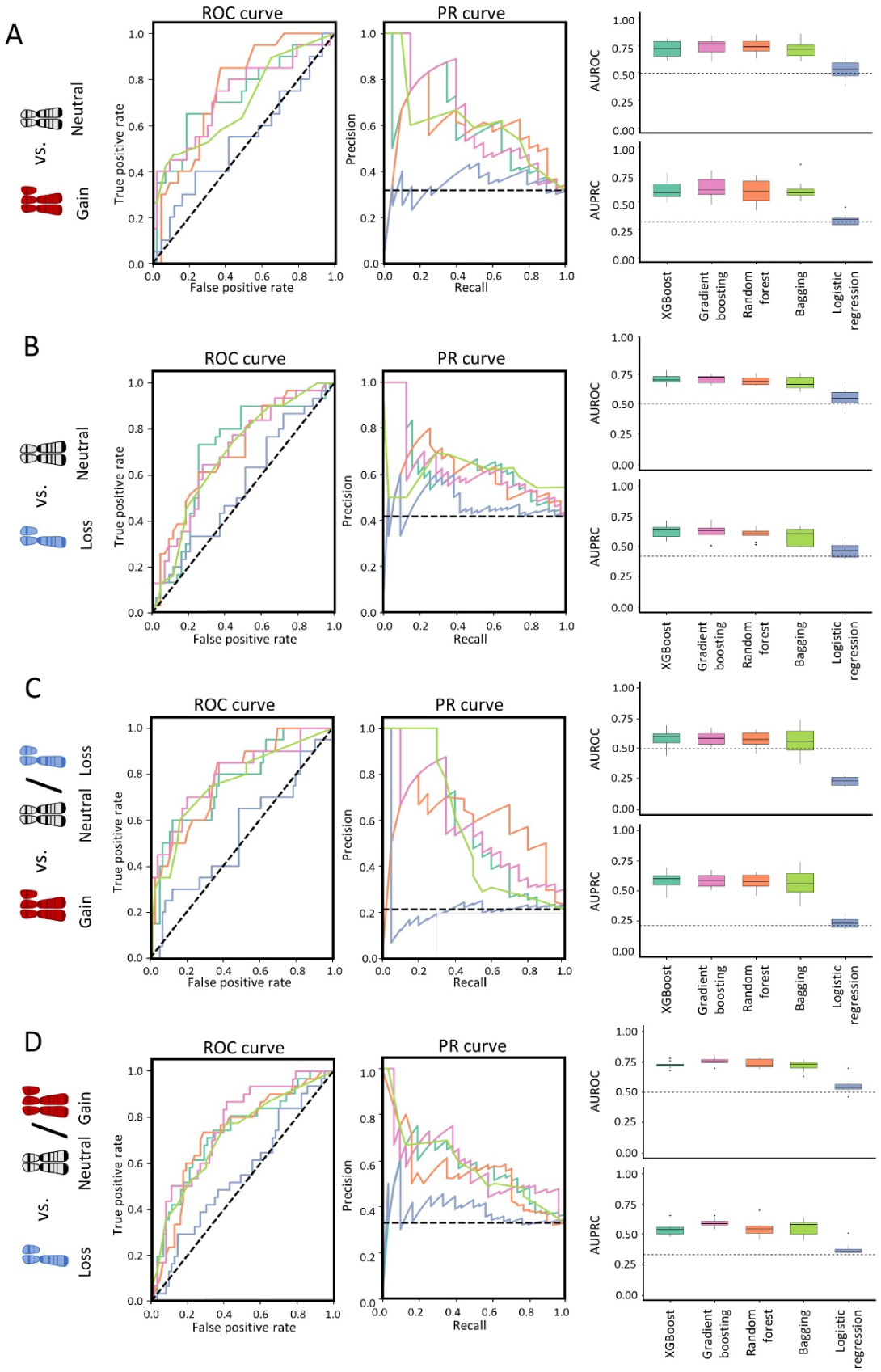
Performance of different ML methods for predicting aneuploidy in cancer. Performance in 10-fold cross-validation of each method was measured by calculating auROC and auPRC. Curves represent the median auROC and auPRC of each method. Boxplots represent the results of auROC and auPRC of all 10 folds. A. Classification between a gain of an arm versus the neutral arms. Gradient boosting method performed best in terms of the auROC and auPRC, and was henceforth used to model chromosome-arm gains. B. Models were applied to classify loss of chromosome-arms versus neutral chromosome-arms. XGBoost performed best in terms of the auROC and auPRC, and was henceforth used to model chromosome-arm loss. C. Models were applied to classify gain of chromosome-arms versus all other chromosome-arms (neutral or lost). XGBoost performed best in terms of the auPRC, and was henceforth used to model chromosome-arm loss. D. Models were applied to classify loss of chromosome-arms versus all other chromosome-arms (neutral or gained). Gradient boosting method performed best in terms of the auROC and auPRC, and was henceforth used to model chromosome-arm gains.

**Figure S4.**
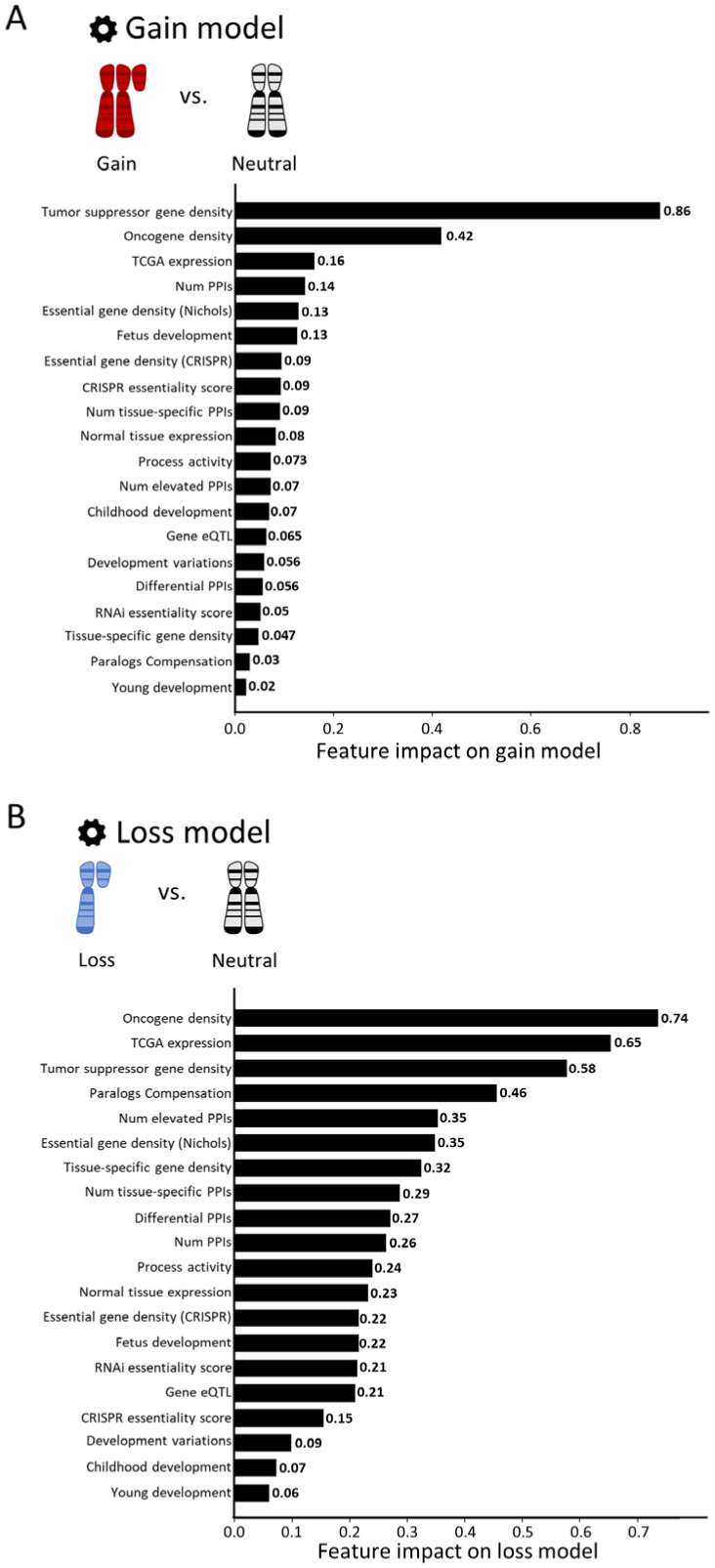
Contribution of all the features to gain and loss models in cancer. The features are ordered from bottom to top by their increased average absolute contribution on the model. A. Gain model. B. Loss model.

**Figure S5.**
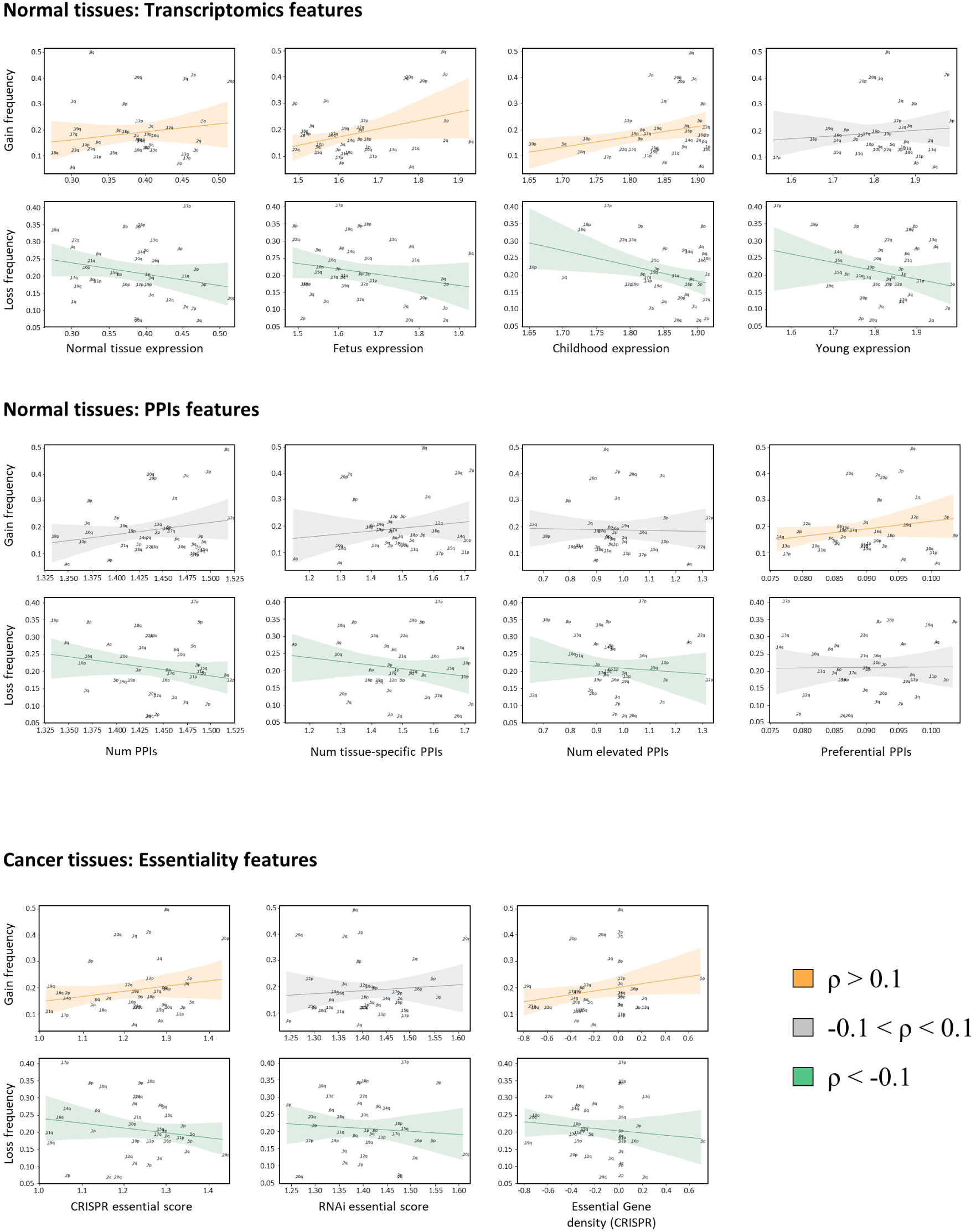
Correlation between different features and the frequencies of chromosome-arms gain and loss. The confidence interval was colored by the Spearman correlation value. Positive correlations (r>0.1) appear in orange, negative correlations (r<-0.1) appear in green, otherwise they appear in grey. A. Transcriptomics features measured in adult (Consortium 2020), fetal (Cardoso-Moreira et al. 2019), child (Cardoso-Moreira et al. 2019) and young subjects (Cardoso-Moreira et al. 2019) were modestly positively correlated with chromosome-arm gain frequency and modestly negatively correlated with chromosome-arm loss. Adult tissues: ρ=0.12, ρ=-0.22 respectively. Fetal tissues: ρ=0.15, ρ=-0.17 respectively. Child tissues: ρ=0.2, ρ=-0.28 respectively. Young tissues: ρ=-0.03, ρ=-0.16 respectively. B. PPI features were generally uncorrelated with chromosome-arm gain frequency and modestly negatively correlated with chromosome-arm loss frequency. Num PPIs: ρ≈0, ρ=-0.18 respectively. Num tissue-specific PPIs: ρ≈0, ρ=-0.13 respectively. Num elevated PPIs: ρ≈0, ρ=-0.13 respectively. Differential PPIs: ρ=0.18, no correlation respectively. C. Gene essentiality features were modestly positively correlated with chromosome-arm gain frequency and modestly negatively correlated with chromosome-arm loss frequency. CRISPR essential score: ρ=0.13, ρ=-0.14 respectively. RNAi essential score: ρ≈0, ρ=-0.12 respectively. Essential gene density (CRISPR): ρ=0.18, ρ=-0.1 respectively.

**Figure S6.**
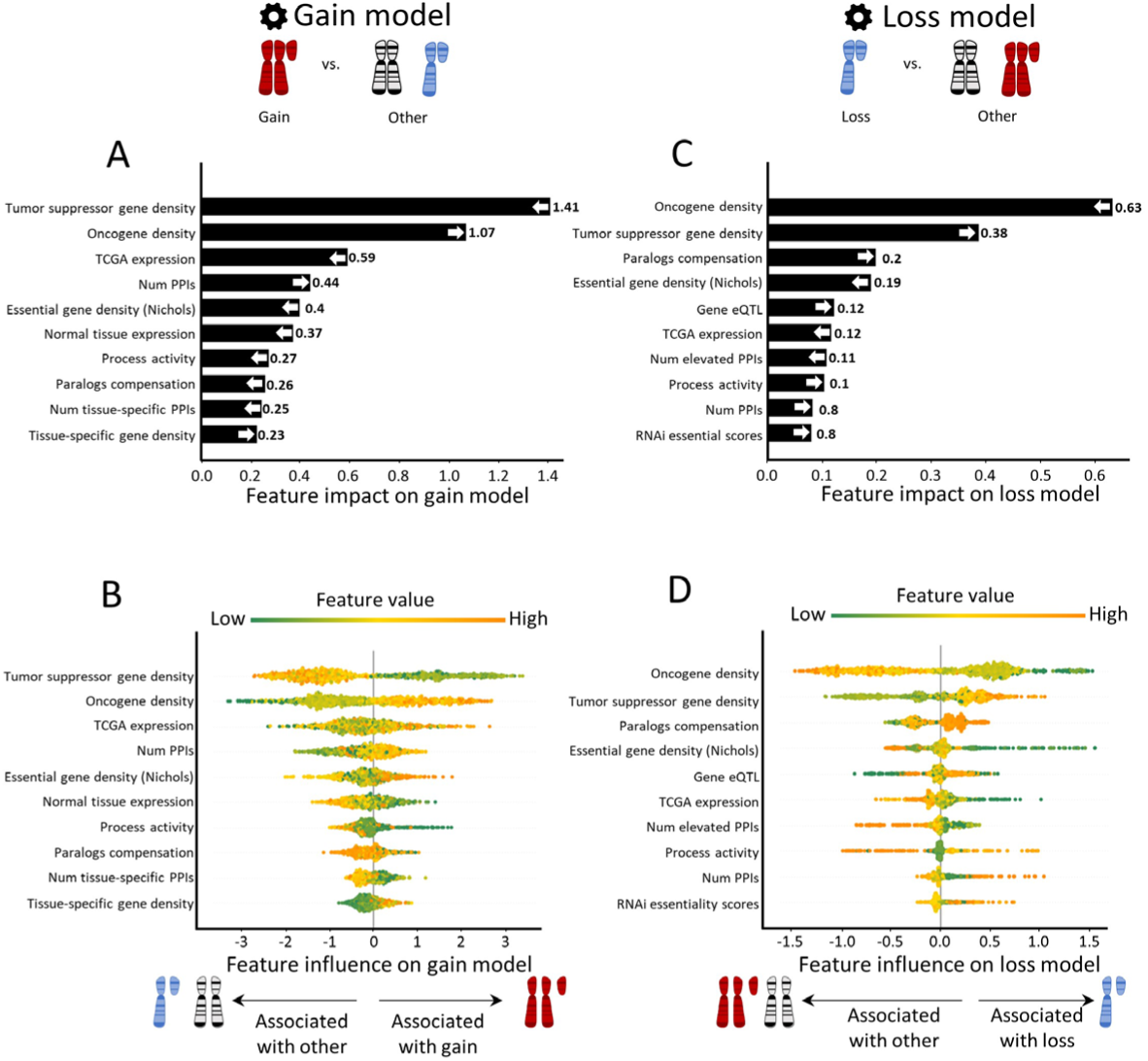
Quantitative view of the 10 topmost contributing features in models that classify gain or loss of chromosome-arms versus all other chromosome-arms. Features are ordered from bottom to top by their increased average absolute contribution on the model. A. The average absolute contribution of each feature to the gain model. B. A detailed view of the contribution of each feature to the gain model. Per feature, each dot represents the contribution per instance of a chromosome-arm and cancer type pair. The dots are spread based on whether they were classified as neutral (left) or gain (right) by the model. Instances are colored by the feature value (green-to-orange scale denotes low-to-high value). The order and directionality of the features generally agree with models of primary tumors (Fig. 2). C. Same as panel A for the loss model. D. Same as panel B for the loss model. The order and directionality of the features generally agree with model of primary tumors (Fig. 2).

**Figure S7:**
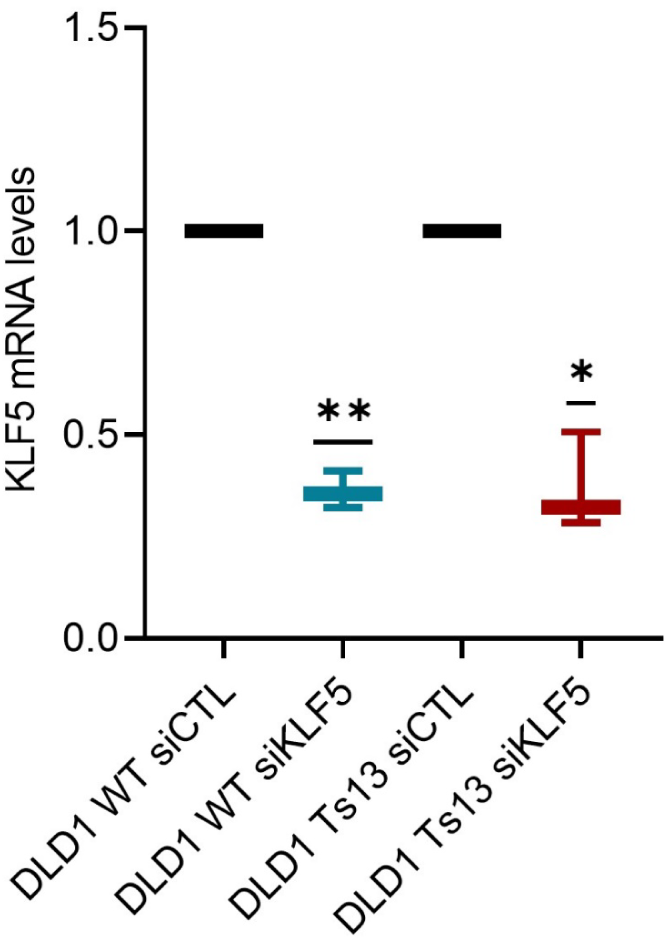
Validation of KLF5 knockdown in DLD1 isogenic cell lines. Comparison of KLF5 mRNA levels between DLD1-WT and DLD1-Ts13 after siRNA treatment against KLF5. Both DLD1-WT and DLD1-Ts13 reached efficient knockdown. n=3 independent experiments. **, p=0.005 and *, p=0.003; One-sample t-test.

**Figure S8.**
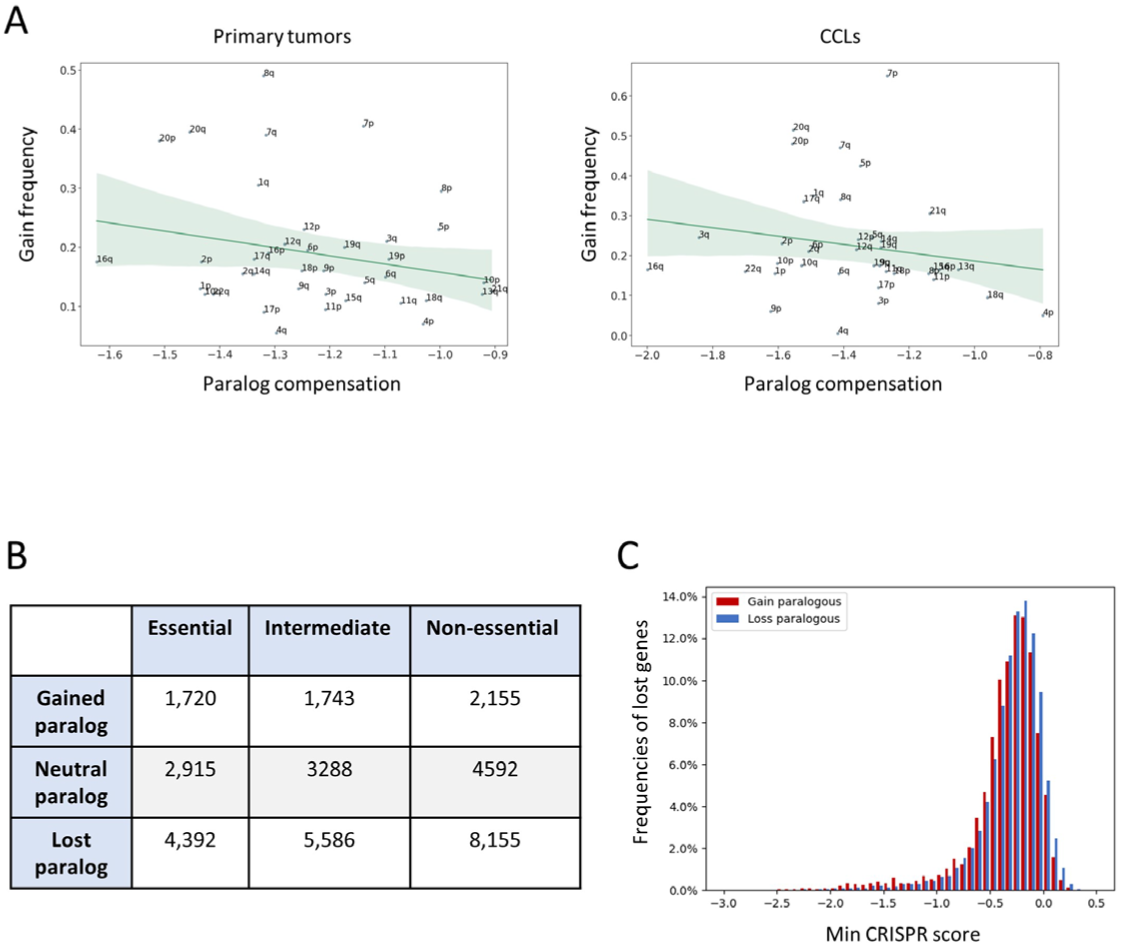
Support for the role of paralog compensation in shaping tissue-specific aneuploidy patterns. A. Correlation between paralogs compensation and chromosome-arm gain frequency in primary tumors (left, ρ=-0.21, adjusted p=0.25) and CCLs (right, ρ=-0.24, adjusted p=0.19, Spearman correlation). Frequently-gained chromosome-arms tend to have low paralog compensation values. B. Analysis of recurrently-lost genes according to their essentiality and the aneuploidy pattern of their paralog (p=2.38e-24, Chi-squared test). C. The distribution of recurrently-lost genes by their essentiality (CRISPR score, (Tsherniak et al. 2017)). Recurrently-lost genes with a frequently-gained paralog tend to be more essential than those with recurrently-lost paralog (p=9.2e-16, KS-test).

## References

Alfieri, F., G. Caravagna, and M. H. Schaefer. 2023. ‘Cancer genomes tolerate deleterious coding mutations through somatic copy number amplifications of wild-type regions’, Nat Commun, 14: 3594.

Barshir, R., I. Hekselman, N. Shemesh, M. Sharon, L. Novack, and E. Yeger-Lotem. 2018. ‘Role of duplicate genes in determining the tissue-selectivity of hereditary diseases’, PLoS Genet, 14: e1007327.

Basha, O., C. M. Argov, R. Artzy, Y. Zoabi, I. Hekselman, L. Alfandari, V. Chalifa-Caspi, and E. Yeger-Lotem. 2020. ‘Differential network analysis of multiple human tissue interactomes highlights tissue-selective processes and genetic disorder genes’, Bioinformatics, 36: 2821–28.

Basha, O., D. Flom, R. Barshir, I. Smoly, S. Tirman, and E. Yeger-Lotem. 2015. ‘MyProteinNet: build up-to-date protein interaction networks for organisms, tissues and user-defined contexts’, Nucleic Acids Res, 43: W258–63.

Ben-David, U., and A. Amon. 2020. ‘Context is everything: aneuploidy in cancer’, Nat Rev Genet, 21: 44–62.

Ben-David, U., G. Ha, Y. Y. Tseng, N. F. Greenwald, C. Oh, J. Shih, J. M. McFarland, B. Wong, J. S. Boehm, R. Beroukhim, and T. R. Golub. 2017. ‘Patient-derived xenografts undergo mouse-specific tumor evolution’, Nat Genet, 49: 1567–75.

Ben-David, U., B. Siranosian, G. Ha, H. Tang, Y. Oren, K. Hinohara, C. A. Strathdee, J. Dempster, N. J. Lyons, R. Burns, A. Nag, G. Kugener, B. Cimini, P. Tsvetkov, Y. E. Maruvka, R. O‘Rourke, A. Garrity, A. A. Tubelli, P. Bandopadhayay, A. Tsherniak, F. Vazquez, B. Wong, C. Birger, M. Ghandi, A. R. Thorner, J. A. Bittker, M. Meyerson, G. Getz, R. Beroukhim, and T. R. Golub. 2018. ‘Genetic and transcriptional evolution alters cancer cell line drug response’, Nature, 560: 325–30.

Benjamini, Yoav, and Yosef Hochberg. 1995. ‘Controlling the false discovery rate: a practical and powerful approach to multiple testing’, Journal of the Royal statistical society: series B (Methodological), 57: 289–300.

Cardoso-Moreira, M., J. Halbert, D. Valloton, B. Velten, C. Chen, Y. Shao, A. Liechti, K. Ascencao, C. Rummel, S. Ovchinnikova, P. V. Mazin, I. Xenarios, K. Harshman, M. Mort, D. N. Cooper, C. Sandi, M. J. Soares, P. G. Ferreira, S. Afonso, M. Carneiro, J. M. A. Turner, J. L. VandeBerg, A. Fallahshahroudi, P. Jensen, R. Behr, S. Lisgo, S. Lindsay, P. Khaitovich, W. Huber, J. Baker, S. Anders, Y. E. Zhang, and H. Kaessmann. 2019. ‘Gene expression across mammalian organ development’, Nature, 571: 505–09.

Chen, C., H. V. Bhalala, H. Qiao, and J. T. Dong. 2002. ‘A possible tumor suppressor role of the KLF5 transcription factor in human breast cancer’, Oncogene, 21: 6567–72.

Chen, Tianqi, and Carlos Guestrin. 2016. “Xgboost: A scalable tree boosting system.” In Proceedings of the 22nd acm sigkdd international conference on knowledge discovery and data mining, 785–94.

Chiu, Yu-Chiao, Siyuan Zheng, Li-Ju Wang, Brian S Iskra, Manjeet K Rao, Peter J Houghton, Yufei Huang, and Yidong Chen. 2021. ‘Predicting and characterizing a cancer dependency map of tumors with deep learning’, Science Advances, 7: eabh1275.

Cohen-Sharir, Y., J. M. McFarland, M. Abdusamad, C. Marquis, S. V. Bernhard, M. Kazachkova, H. Tang, M. R. Ippolito, K. Laue, J. Zerbib, H. L. H. Malaby, A. Jones, L. M. Stautmeister, I. Bockaj, R. Wardenaar, N. Lyons, A. Nagaraja, A. J. Bass, D. C. J. Spierings, F. Foijer, R. Beroukhim, S. Santaguida, T. R. Golub, J. Stumpff, Z. Storchova, and U. Ben-David. 2021. ‘Aneuploidy renders cancer cells vulnerable to mitotic checkpoint inhibition’, Nature, 590: 486–91.

Consortium, GTEx. 2020. ‘The GTEx Consortium atlas of genetic regulatory effects across human tissues’, Science, 369: 1318–30.

Cunningham, F., J. E. Allen, J. Allen, J. Alvarez-Jarreta, M. R. Amode, I. M. Armean, O. Austine-Orimoloye, A. G. Azov, I. Barnes, R. Bennett, A. Berry, J. Bhai, A. Bignell, K. Billis, S. Boddu, L. Brooks, M. Charkhchi, C. Cummins, L. Da Rin Fioretto, C. Davidson, K. Dodiya, S. Donaldson, B. El Houdaigui, T. El Naboulsi, R. Fatima, C. G. Giron, T. Genez, J. G. Martinez, C. Guijarro-Clarke, A. Gymer, M. Hardy, Z. Hollis, T. Hourlier, T. Hunt, T. Juettemann, V. Kaikala, M. Kay, I. Lavidas, T. Le, D. Lemos, J. C. Marugan, S. Mohanan, A. Mushtaq, M. Naven, D. N. Ogeh, A. Parker, A. Parton, M. Perry, I. Pilizota, I. Prosovetskaia, M. P. Sakthivel, A. I. A. Salam, B. M. Schmitt, H. Schuilenburg, D. Sheppard, J. G. Perez-Silva, W. Stark, E. Steed, K. Sutinen, R. Sukumaran, D. Sumathipala, M. M. Suner, M. Szpak, A. Thormann, F. F. Tricomi, D. Urbina-Gomez, A. Veidenberg, T. A. Walsh, B. Walts, N. Willhoft, A. Winterbottom, E. Wass, M. Chakiachvili, B. Flint, A. Frankish, S. Giorgetti, L. Haggerty, S. E. Hunt, I. Isley GR, J. E. Loveland, F. J. Martin, B. Moore, J. M. Mudge, M. Muffato, E. Perry, M. Ruffier, J. Tate, D. Thybert, S. J. Trevanion, S. Dyer, P. W. Harrison, K. L. Howe, A. D. Yates, D. R. Zerbino, and P. Flicek. 2022. ‘Ensembl 2022’, Nucleic Acids Res, 50: D988–D95.

Davoli, T., A. W. Xu, K. E. Mengwasser, L. M. Sack, J. C. Yoon, P. J. Park, and S. J. Elledge. 2013. ‘Cumulative haploinsufficiency and triplosensitivity drive aneuploidy patterns and shape the cancer genome’, Cell, 155: 948–62.

Goldman, M. J., B. Craft, M. Hastie, K. Repecka, F. McDade, A. Kamath, A. Banerjee, Y. Luo, D. Rogers, A. N. Brooks, J. Zhu, and D. Haussler. 2020. ‘Visualizing and interpreting cancer genomics data via the Xena platform’, Nat Biotechnol, 38: 675–78.

Greene, C. S., A. Krishnan, A. K. Wong, E. Ricciotti, R. A. Zelaya, D. S. Himmelstein, R. Zhang, B. M. Hartmann, E. Zaslavsky, S. C. Sealfon, D. I. Chasman, G. A. FitzGerald, K. Dolinski, T. Grosser, and O. G. Troyanskaya. 2015. ‘Understanding multicellular function and disease with human tissue-specific networks’, Nat Genet, 47: 569–76.

Han, Y., J. Yang, X. Qian, W. C. Cheng, S. H. Liu, X. Hua, L. Zhou, Y. Yang, Q. Wu, P. Liu, and Y. Lu. 2019. ‘DriverML: a machine learning algorithm for identifying driver genes in cancer sequencing studies’, Nucleic Acids Res, 47: e45.

Hua, Jianping, Zixiang Xiong, James Lowey, Edward Suh, and Edward R Dougherty. 2005. ‘Optimal number of features as a function of sample size for various classification rules’, Bioinformatics, 21: 1509–15.

Ito, T., M. J. Young, R. Li, S. Jain, A. Wernitznig, J. M. Krill-Burger, C. T. Lemke, D. Monducci, D. J. Rodriguez, L. Chang, S. Dutta, D. Pal, B. R. Paolella, M. V. Rothberg, D. E. Root, C. M. Johannessen, L. Parida, G. Getz, F. Vazquez, J. G. Doench, M. Zamanighomi, and W. R. Sellers. 2021. ‘Paralog knockout profiling identifies DUSP4 and DUSP6 as a digenic dependence in MAPK pathway-driven cancers’, Nat Genet, 53: 1664–72.

Jubran, J., I. Hekselman, L. Novack, and E. Yeger-Lotem. 2020. ‘Dosage-sensitive molecular mechanisms are associated with the tissue-specificity of traits and diseases’, Comput Struct Biotechnol J, 18: 4024–32.

Kegel, Barbara De, and Colm J. Ryan. 2022. ‘Paralog dispensability shapes homozygous deletion patterns in tumor genomes’, bioRxiv: 2022.06.20.496722.

Kingsford, C., and S. L. Salzberg. 2008. ‘What are decision trees?’, *Nat Biotechnol*, 26: 1011-3. Kotsiantis, Sotiris B. 2013. ‘Decision trees: a recent overview‘, Artificial Intelligence Review, 39: 261–83.

Liu, Y., C. Chen, Z. Xu, C. Scuoppo, C. D. Rillahan, J. Gao, B. Spitzer, B. Bosbach, E. R. Kastenhuber, T. Baslan, S. Ackermann, L. Cheng, Q. Wang, T. Niu, N. Schultz, R. L. Levine, A. A. Mills, and S. W. Lowe. 2016. ‘Deletions linked to TP53 loss drive cancer through p53-independent mechanisms’, Nature, 531: 471–75.

Lundberg, S. M., G. Erion, H. Chen, A. DeGrave, J. M. Prutkin, B. Nair, R. Katz, J. Himmelfarb, N. Bansal, and S. I. Lee. 2020. ‘From Local Explanations to Global Understanding with Explainable AI for Trees’, Nat Mach Intell, 2: 56–67.

Lundberg, Scott M, and Su-In Lee. 2017. ‘A unified approach to interpreting model predictions’, Advances in neural information processing systems, 30.

Luo, P., Y. Ding, X. Lei, and F. X. Wu. 2019. ‘deepDriver: Predicting Cancer Driver Genes Based on Somatic Mutations Using Deep Convolutional Neural Networks’, Front Genet, 10: 13.

Luo, Y., and C. Chen. 2021. ‘The roles and regulation of the KLF5 transcription factor in cancers’, Cancer Sci, 112: 2097–117.

Ma, Jian-Bin, Ji-Yu Bai, Hai-Bao Zhang, Jing Jia, Qi Shi, Chao Yang, Xinyang Wang, Dalin He, and Peng Guo. 2020. ‘KLF5 inhibits STAT3 activity and tumor metastasis in prostate cancer by suppressing IGF1 transcription cooperatively with HDAC1’, Cell death & disease, 11: 466.

Martincorena, I., K. M. Raine, M. Gerstung, K. J. Dawson, K. Haase, P. Van Loo, H. Davies, M. R. Stratton, and P. J. Campbell. 2017. ‘Universal Patterns of Selection in Cancer and Somatic Tissues’, Cell, 171: 1029–41 e21.

McConnell, B. B., A. B. Bialkowska, M. O. Nandan, A. M. Ghaleb, F. J. Gordon, and V. W. Yang. 2009. ‘Haploinsufficiency of Kruppel-like factor 5 rescues the tumor-initiating effect of the Apc(Min) mutation in the intestine’, Cancer Res, 69: 4125–33.

McFarland, J. M., Z. V. Ho, G. Kugener, J. M. Dempster, P. G. Montgomery, J. G. Bryan, J. M. Krill-Burger, T. M. Green, F. Vazquez, J. S. Boehm, T. R. Golub, W. C. Hahn, D. E. Root, and A. Tsherniak. 2018. ‘Improved estimation of cancer dependencies from large-scale RNAi screens using model-based normalization and data integration’, Nat Commun, 9: 4610.

Mermel, C. H., S. E. Schumacher, B. Hill, M. L. Meyerson, R. Beroukhim, and G. Getz. 2011. ‘GISTIC2.0 facilitates sensitive and confident localization of the targets of focal somatic copy-number alteration in human cancers’, Genome Biol, 12: R41.

Mostavi, M., Y. C. Chiu, Y. Chen, and Y. Huang. 2021. ‘CancerSiamese: one-shot learning for predicting primary and metastatic tumor types unseen during model training’, BMC Bioinformatics, 22: 244.

Nichols, C. A., W. J. Gibson, M. S. Brown, J. A. Kosmicki, J. P. Busanovich, H. Wei, L. M. Urbanski, N. Curimjee, A. C. Berger, G. F. Gao, A. D. Cherniack, S. Dhe-Paganon, B. R. Paolella, and R. Beroukhim. 2020. ‘Loss of heterozygosity of essential genes represents a widespread class of potential cancer vulnerabilities’, Nat Commun, 11: 2517.

Patkar, S., K. Heselmeyer-Haddad, N. Auslander, D. Hirsch, J. Camps, D. Bronder, M. Brown, W. D. Chen, R. Lokanga, D. Wangsa, D. Wangsa, Y. Hu, A. Lischka, R. Braun, G. Emons, B. M. Ghadimi, J. Gaedcke, M. Grade, C. Montagna, Y. Lazebnik, M. J. Difilippantonio, J. K. Habermann, G. Auer, E. Ruppin, and T. Ried. 2021. ‘Hard wiring of normal tissue-specific chromosome-wide gene expression levels is an additional factor driving cancer type-specific aneuploidies’, Genome Med, 13: 93.

Pedregosa, Fabian, Gaël Varoquaux, Alexandre Gramfort, Vincent Michel, Bertrand Thirion, Olivier Grisel, Mathieu Blondel, Peter Prettenhofer, Ron Weiss, and Vincent Dubourg. 2011. ‘Scikit-learn: Machine learning in Python‘, the Journal of machine Learning research, 12: 2825–30.

Prasad, K., M. Bloomfield, H. Levi, K. Keuper, S. V. Bernhard, N. C. Baudoin, G. Leor, Y. Eliezer, M. Giam, C. K. Wong, G. Rancati, Z. Storchova, D. Cimini, and U. Ben-David. 2022. ‘Whole-Genome Duplication Shapes the Aneuploidy Landscape of Human Cancers’, Cancer Res, 82: 1736–52.

Ramirez, R., Y. C. Chiu, A. Hererra, M. Mostavi, J. Ramirez, Y. Chen, Y. Huang, and Y. F. Jin. 2020. ‘Classification of Cancer Types Using Graph Convolutional Neural Networks’, Front Phys, 8.

Rodriguez-Perez, R., and J. Bajorath. 2020. ‘Interpretation of Compound Activity Predictions from Complex Machine Learning Models Using Local Approximations and Shapley Values’, J Med Chem, 63: 8761–77.

Rutledge, S. D., T. A. Douglas, J. M. Nicholson, M. Vila-Casadesus, C. L. Kantzler, D. Wangsa, M. Barroso-Vilares, S. D. Kale, E. Logarinho, and D. Cimini. 2016. ‘Selective advantage of trisomic human cells cultured in non-standard conditions’, Sci Rep, 6: 22828.

Sack, L. M., T. Davoli, M. Z. Li, Y. Li, Q. Xu, K. Naxerova, E. C. Wooten, R. J. Bernardi, T. D. Martin, T. Chen, Y. Leng, A. C. Liang, K. A. Scorsone, T. F. Westbrook, K. K. Wong, and S. J. Elledge. 2018. ‘Profound Tissue Specificity in Proliferation Control Underlies Cancer Drivers and Aneuploidy Patterns’, Cell, 173: 499–514 e23.

Sharon, M., E. Vinogradov, C. M. Argov, O. Lazarescu, Y. Zoabi, I. Hekselman, and E. Yeger-Lotem. 2022. ‘The differential activity of biological processes in tissues and cell subsets can illuminate disease-related processes and cell-type identities’, Bioinformatics, 38: 1584–92.

Sheltzer, J. M., and A. Amon. 2011. ‘The aneuploidy paradox: costs and benefits of an incorrect karyotype’, Trends Genet, 27: 446–53.

Shih, J., S. Sarmashghi, N. Zhakula-Kostadinova, S. Zhang, Y. Georgis, S. H. Hoyt, M. S. Cuoco, G. F. Gao, L. F. Spurr, A. C. Berger, G. Ha, V. Rendo, H. Shen, M. Meyerson, A. D. Cherniack, A. M. Taylor, and R. Beroukhim. 2023. ‘Cancer aneuploidies are shaped primarily by effects on tumour fitness’, Nature, https://doi.org/10.1038/s41586-023-06266-3.

Shukla, A., T. H. M. Nguyen, S. B. Moka, J. J. Ellis, J. P. Grady, H. Oey, A. S. Cristino, K. K. Khanna, D. P. Kroese, L. Krause, E. Dray, J. L. Fink, and P. H. G. Duijf. 2020. ‘Chromosome arm aneuploidies shape tumour evolution and drug response’, Nat Commun, 11: 449.

Simonovsky, E., M. Sharon, M. Ziv, O. Mauer, I. Hekselman, J. Jubran, E. Vinogradov, C. M. Argov, O. Basha, L. Kerber, Y. Yogev, A. V. Segre, H. K. Im, G. TEx Consortium, O. Birk, L. Rokach, and E. Yeger-Lotem. 2023. ‘Predicting molecular mechanisms of hereditary diseases by using their tissue-selective manifestation’, Mol Syst Biol: e11407.

Sonawane, Abhijeet Rajendra, John Platig, Maud Fagny, Cho-Yi Chen, Joseph Nathaniel Paulson, Camila Miranda Lopes-Ramos, Dawn Lisa DeMeo, John Quackenbush, Kimberly Glass, and Marieke Lydia Kuijjer. 2017. ‘Understanding tissue-specific gene regulation’, Cell reports, 21: 1077–88.

Taylor, A. M., J. Shih, G. Ha, G. F. Gao, X. Zhang, A. C. Berger, S. E. Schumacher, C. Wang, H. Hu, J. Liu, A. J. Lazar, Network Cancer Genome Atlas Research, A. D. Cherniack, R. Beroukhim, and M. Meyerson. 2018. ‘Genomic and Functional Approaches to Understanding Cancer Aneuploidy’, Cancer Cell, 33: 676–89 e3.

Tsherniak, A., F. Vazquez, P. G. Montgomery, B. A. Weir, G. Kryukov, G. S. Cowley, S. Gill, W. F. Harrington, S. Pantel, J. M. Krill-Burger, R. M. Meyers, L. Ali, A. Goodale, Y. Lee, G. Jiang, J. Hsiao, W. F. J. Gerath, S. Howell, E. Merkel, M. Ghandi, L. A. Garraway, D. E. Root, T. R. Golub, J. S. Boehm, and W. C. Hahn. 2017. ‘Defining a Cancer Dependency Map’, Cell, 170: 564–76 e16.

Vasudevan, A., P. S. Baruah, J. C. Smith, Z. Wang, N. M. Sayles, P. Andrews, J. Kendall, J. Leu, N. K. Chunduri, D. Levy, M. Wigler, Z. Storchova, and J. M. Sheltzer. 2020. ‘Single-Chromosomal Gains Can Function as Metastasis Suppressors and Promoters in Colon Cancer’, Dev Cell, 52: 413–28 e6.

Wang, T., K. Birsoy, N. W. Hughes, K. M. Krupczak, Y. Post, J. J. Wei, E. S. Lander, and D. M. Sabatini. 2015. ‘Identification and characterization of essential genes in the human genome’, Science, 350: 1096–101.

Zapata, Luis, Oriol Pich, Luis Serrano, Fyodor A Kondrashov, Stephan Ossowski, and Martin H Schaefer. 2018. ‘Negative selection in tumor genome evolution acts on essential cellular functions and the immunopeptidome’, Genome biology, 19: 1–17.

Zhou, X. P., Y. J. Li, K. Hoang-Xuan, P. Laurent-Puig, K. Mokhtari, M. Longy, M. Sanson, J. Y. Delattre, G. Thomas, and R. Hamelin. 1999. ‘Mutational analysis of the PTEN gene in gliomas: molecular and pathological correlations’, Int J Cancer, 84: 150–4.

Zitnik, M., F. Nguyen, B. Wang, J. Leskovec, A. Goldenberg, and M. M. Hoffman. 2019. ‘Machine Learning for Integrating Data in Biology and Medicine: Principles, Practice, and Opportunities’, Inf Fusion, 50: 71–91.

